# Multimodal GiaC-phenomic approach for microbiome-tree system profiling

**DOI:** 10.1101/2025.11.26.690777

**Authors:** Maksymilian Chmielewski, Jakub Kuncewicz, Władysław Polcyn

## Abstract

We constructed a computational methodology to assess health of plant-microbiome system through microbiome structure modelling combined with plant remote sensing. As a test dataset, we selected soil mycobiome and morphometry of *Tilia cordata* in nursery and forest sites. Our method is also applicable on forest or regional scale.

Microbiome part called GiaC (**G**u**i**lds **a**nd o**C**currences) combines taxonomic and trophic composition as well as species co-occurrence modelled with advanced graph methods. We complemented state-of-the-art approaches with novel ones for visualisations, species filtering (Flexible99) and graph transformation modelling species clusters (ClusterCollapse). Flexible99 is a method that adjusts the species abundance cut-off to each sample set and removes rare species. ClusterCollapse generalises co-occurrence networks to species clusters by edge contraction and serves as an implicit homogeneity test.

To assess biomass of the seedlings we used low-cost and field-adopted morphometric and manual measurements. Top and side tree images, acquired with handheld RGB camera, were analysed using colour segmentation and pixel count based methods. Parameters, such as crown size, shape, area and pigment content, number of leaves, branch length and foliage density, allowed the seedlings to be classified into three different vitality groups.

Presented multimodal approach was capable to differentiate and characterize distinct best, suboptimal or critical states of microbiome-host system, both on microbial and plant side. Our results show that more stable fungal co-occurrence patterns should be attributed to the plant set of the best growth. In contrast, more chaotic patterns can be considered non-optimal for plant-mycobiome cooperation.

## 1. Introduction

Macroorganisms, including plants, are inherently associated with complex microbiomes through physical and chemical interactions, forming holobiont systems. To understand these systems, it is crucial to study both microbial and plant behaviours (Cordovez et al. 2019; Hassani et al. 2018; Mesny et al. 2023). This can be effectively realized through a multifaceted workflow using microbiome focused metagenomics and plants’ remote sensing (Beatty et al. 2021; Sørensen et al. 2025).

Metagenomic approach provides massive amounts of data about microbiota, although requires advanced computational methods on each analytical step (Bharti et al. 2021; Lucaciu et al. 2019). Despite high-throughput efficiency metagenomics generate some level of statistical uncertainty arising from limited volume of samples and spatial dispersion of sampling but also from limits of used sequencing method. Obtaining complete information on microbial communities is difficult also due to compounds interfering with their isolation that require specialised isolation techniques and very high species numbers (Lamb et al. 2019; Tedersoo et al. 2022).

High number of species in a plant rhizosphere microbiome requires proper analytical methods, such as graph theory, to reason about their relationships I (Barberán et al. 2012). Graphs represent objects and their relationships. Graph analysis, albeit computationally challenging, can infer ecological relationships from species occurrence data using methods designed for sparse data as many tools assume biological networks are sparse (Kurtz et al. 2015). Sparse data analysis methods assume that data is composed in a significant proportion of missing, irrelevant, or zero-valued elements (Ye and Liu 2012). These methods make them suitable for metagenomics, as most of the species occur in less than all samples and metagenomics do not represent real occurrence values. Barcode metagenomics provide direct information only on species composition. However, using literature knowledge or genome analysis, it is possible to translate species data into functional diversity by assigning them an ecological role (Nguyen et al. 2016; Põlme et al. 2020, Tanunchai et al. 2023).

Functional relationships can be considered both at the level of species-species interactions but also in the form of group interactions, grouping different species into clusters. Clusters can be constructed to represent similar functions, similar occurrence patterns or genetic markers.

To analyse the plant side of the holobiont remote plant phenotyping is well suited to evaluate the influence that microbiome fungi enact upon macroorganisms they cohabitate with. These methods integrate non-invasive, high-throughput analyses vastly improving data quality providing insights into factors affecting plant health, their changes over time in yield or disruptions in their phenological phases (Abebe et al., 2023; Humplík et al., 2015). Recent advancements in miniaturization and modularity have enabled common commercial devices, such as smartphones, to reliably function as cheap, easy to use and readily available RBG sensors and serve as competitive alternative when more extensive solutions are not warranted (Liu et al. 2021; Yadav and Yadav, K. 2025).

In this paper we present a proof of concept of a methodology that combines metagenomic assessment of fungal rhizobiome and remote sensing of plant condition. That allows us to analyse plant-microbiome system as a single, holistic entity and confirm whether our methodology is sensitive enough to detect variation in the microbiome structure associated with different plant vitality.

## 2. Methodology description

### Test data and hypotheses

It is crucial to address a key feature of our methodology: the absence of any control sets. Our methodology does not measure any parameters or features as deviations from some norm but uses absolute values. This allows it to be used both to compare multiple sets as well as to simply characterise singular set. One could go and study the quirkiest environment they can find and still would get full results, with no need to define control for it.

We focused ourselves to barcoding of rhizosphere fungi data on metagenomic side and RGB based plant morphometry on remote sensing side. That allowed us to test our concepts on simple, but representative data sets. The development of methodology was based on 26 small-leaved lime (*Tilia cordata*) seedlings (1.5 m high), previously grown as the same nursery batch, categorized into two site-related groups: one still grown in nursery pots – 11 trees (“Pots” group), and one planted on a forest plots (Darzybór forest complex, Poznań, Poland) - 15 trees (“Forest” group). Through a preliminary morphometric assessment, we identified three distinct states of seedling vitality in both sites, and categorized them by differences in growth delays. “Normal” seedlings exhibited the best morphological parameters. “Retarded” were severely stunted seedlings having small residual leafage with severe up to sublethal growth impairment. “Delayed” seedlings showed mixed growth retardation in relation to Normal group but significantly more healthy than Retarded group, accompanied by smaller number of leaves or branches. In Pots there where 5 Normal, 3 Retarded and 3 Delayed. In Forest there were 8 Normal, 4 Delayed and 3 Retarded. We sampled the soil from under each condition differently giving us 10 Normal, 9 Delayed and 9 Retarded soil samples in Pots with 8 Normal, 8 Delayed and 9 Retarded soil samples in Forest. This way we managed to balance out uneven number of samples in each condition.

We outlined three hypotheses to test the extent to which our methodology is sensitive in detecting variations in the microbiome’s structure, as the taxonomic and functional composition as well as relationships between species.

Hypothesis 1: Different plant conditions are accompanied by different mycobiome structure.
Hypothesis 2: Pots and Forest plants of corresponding condition is accompanied by a similar mycobiome structure.
Hypothesis 3: The structure of the mycobiome accompanying the intermediate plant condition has characteristics halfway between those of the best and worst plant condition.

### 2.1. GiaC

Here we present 3 Stages of GiaC (**G**u**i**lds **a**nd o**C**currences) procedure. It connects taxonomy and function of detected species and analyses fungal composition on both of those levels. Thanks to that combined analysis, it characterises microbiome structure as the taxonomic composition, functional composition as well as relationships between species. It combines methods aiming to analyse quantitative composition (e.g. Venn diagram on number of species or reads), qualitative composition (e.g. hierarchical clustering of taxonomic composition) and co-occurrence networks. GiaC combines all those different methods to create a comprehensive picture of metagenomic data from multiple analytical angles. While here it is stated as a method to study fungal microbiome, is can be applied it to any other metagenomic data if methods in Stage I are adjusted to given data.

#### 2.1.1. Stage I – Taxonomic and trophic assignment

Stage I can be simply defined as noise removal and data preprocessing. In metagenomics, noise is a combination of bad sequencing reads, uncertain taxonomic assignment and removal of unimportant species. Due to fast pace of development of metagenomic tools and resources, anyone trying to implement this methodology is advised to research best tools and databases for each analytical step at the time of implementing it. We have primarily used popular tools with default flags and parameters to prove that even the default way gives informative results.

##### Raw data and quality control

The beginning of the procedure consists of getting eDNA sequenced after its isolation from biological material. After the sequencing is done, filtering, demultiplexing and general quality control of raw sequencing reads needs to be performed. The tools and parameters used depend on sequencing technology and machine used, as they change characteristics like sequencing depths or read length.

##### Barcode sequence grouping

After the data is clean and separated by samples, ASV table needs to be created. Tools and methods at this stage must be carefully picked as the way ASV sequences are created has influence on later stages of analysis. This and the previous step aim to remove or correct noise from erroneous sequencing reads.

##### Taxonomic and functional assignment

ASV sequences are used for taxonomic identification. We advise to try few different tools for taxonomic assignment as it may show what species or genus are hard to identify or what ASVs are uncertain. After the taxonomic identification next step is to identify what functional groups the species fall into. For fungal species it can be done using databases such as FungalTraits (Põlme et al. 2020) and FUNGuild (Nguyen et al. 2016) or one can attempt analysis of genomes for available species. For taxonomic assignment removal of noise is based on statistics such as e-value and similar. It is very hard to conceive any such parameters for functional assignment based on databases, but it might be possible for genomic analyses. Due to high number of mycorrhiza subtypes or other trophic types such a pathogen, we decided to group similar lifestyles into more general guilds (Myc - Mycorhizae, Sap - Saprotroph, End - Endophyte, Pat – fungal or plant targeted pathogens, Npa –not plant specific and ENA – Ecologically Not Assigned). For details of guild categorisation of FungalTraits terms, see ‘Data Availability’ section.

##### Species filtering - Flexible99

After all those steps are performed, it is recommended to use some species filtering method to focus on species relevant to studied site. Proper filtering can lead to more reliable, replicable and comparable results in downstream analysis (Cao et al. 2021). We created abundance-based method called Flexible99 (F99) with flexible cut-off point. We quantify abundance as number of reads assigned to a specific species. F99 keeps the species that together make up 99% of all reads in a given sample set. This method first calculates what numerical value 99% of the summed abundances for that sample set would be. Then it sorts all species based on their abundance in decreasing order. And the final step is going through that sorted list of species starting with the most abundant and adding up their numbers. When the 99% point is reached, algorithm stops. It later removes species that pass abundance filter but occur in a single sample only and returns final filtered set. This is the last part of the noise removal. See chapter ‘Method effectiveness’ for justification of this method. Example results are presented on Figure 1.

**Figure 1.**
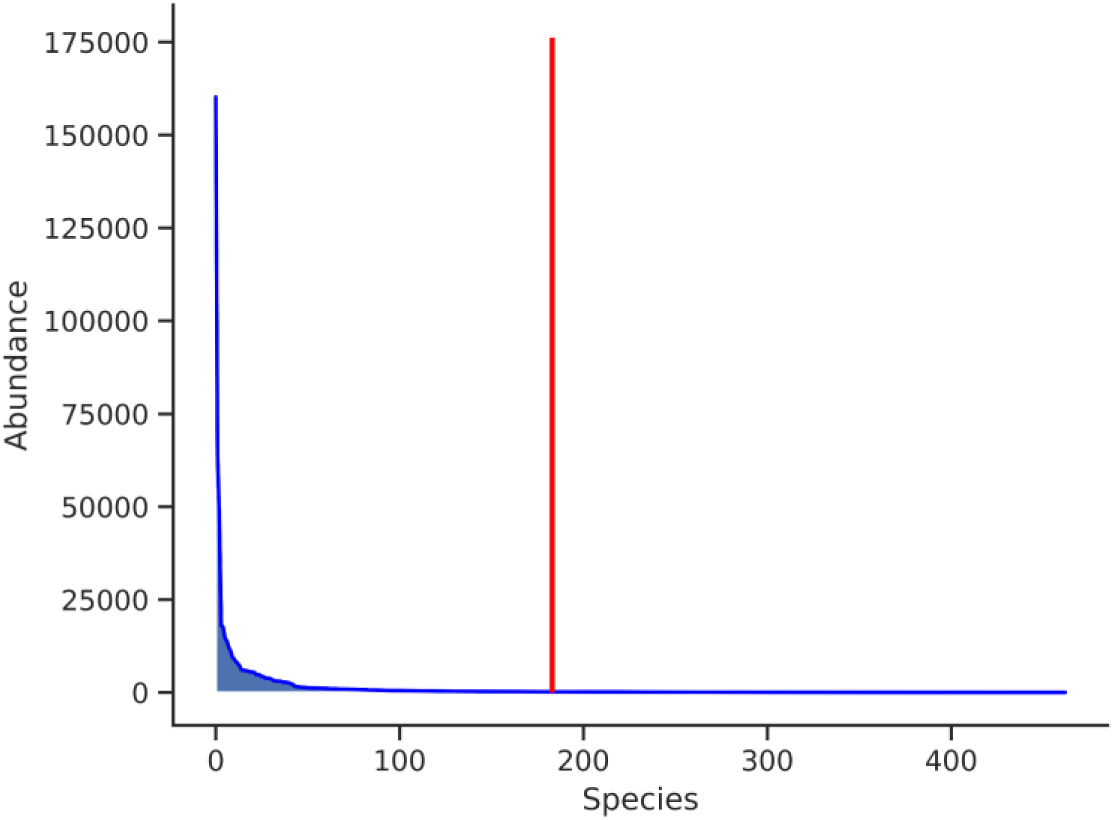
Example of F99 results based on filtering species inhabiting rhizosphere of Normal Pots variant. Blue line represents abundance level of species sorted from the most abundant. Vertical red line notes the cut-off value. Shown here are species with summary abundance of at least 1 in this sample set.

##### Sankey diagram

The last step of Stage I is generation of flow diagram also known as Sankey diagram (Kennedy and Sankey 1898). This is a type of visualisation that shows how some number of some objects changes during the process. It stands out from other flow chart types as it includes graphical representation of amount of the object on each stage, not only a numerical label. In our case it is used to show how many reads pass subsequent steps of filtering. We have created a layout that shows all stages of filtering, taxonomic assignment and functional assignment, as well as F99 filtering step (Fig. 2.).

**Figure 2.**
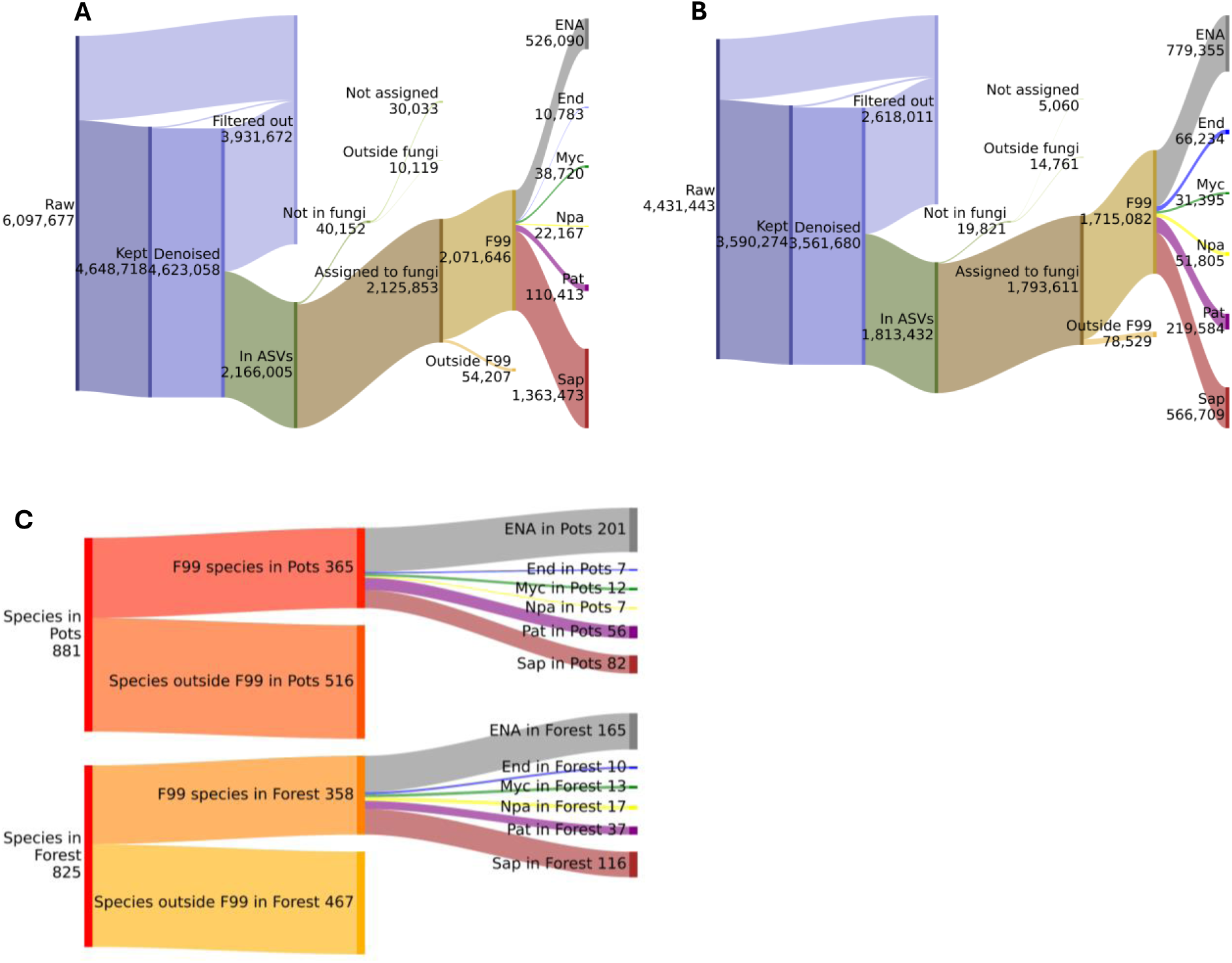
Sankey diagram template with flow of reads shown for Forest (A) and Pots (B) sites and species flow (C) shown for both sites. This presents how many reads pass subsequent GiaC steps and number of species assigned to each trophic guild in a concise and simple manner.

#### 2.1.2. Stage II – Microbiome composition analysis

The composition analysis is split into 2 parts: taxonomic and functional. Most of tools we list as appropriate for this stage are good for both types of composition. Sankey flow charts already provide a summary of functional composition both in species number and abundance value, but they don’t show taxonomic composition nor make it easy to compare several datasets. For those purposes, we have selected an assortment of several well-known techniques such as Venn diagrams and clustermaps. Here we describe how they can be used and state major limitations for each visualisation if present.

##### Venn diagram

Simple way to check roughly how similar sample sets are is Venn diagram. They can be used to compare taxonomic or functional composition. Three main issues of Venn diagram are binary existence used for comparison, no representation of internal variability of sample set and fixed sets specified as input.

##### Clustermap

Those shortcomings can be compensated by technique called clustermap. It is a modification of a heatmap with clustering of columns and rows performed and shown as a dendrogram. Clustermap is a method to analyse internal variability of a given dataset not how two sets compare. To analyse two disjoined sets, they need to be combined together. We advise to set samples as columns and rows as species; as well as toggling the clustering of rows and columns on and clustering method set to Ward’s method (Müllner, D. 2011). Ward’s method combines two samples or clusters thereof that lead to a new, biggest cluster with the minimal internal variance. Clustermap will not only take into account existence of a species in a given sample, but it will also their abundance. Proper normalisation method for clustermap is percentile method applied to samples. Percentile method is very good approach for sparse data analysis as it will very clearly mark values that are 0 or > 0.

##### Guild ratios

Last of the issues solved by this stage is simple way to compare multiple functional guild compositions at once. For a comprehensive comparison of species in a single guild, we advise a clustermap. For a more general comparison a simple grouped barplot showing either sum of abundance or number of species in each guild is a suitable choice. Using bars instead of pie charts make it easier to compare inside a sample set or between multiple ones. We advise to always show both absolute and percentage values on two panels. An example barplot can be seen on a Figure 3.

**Figure 3.**
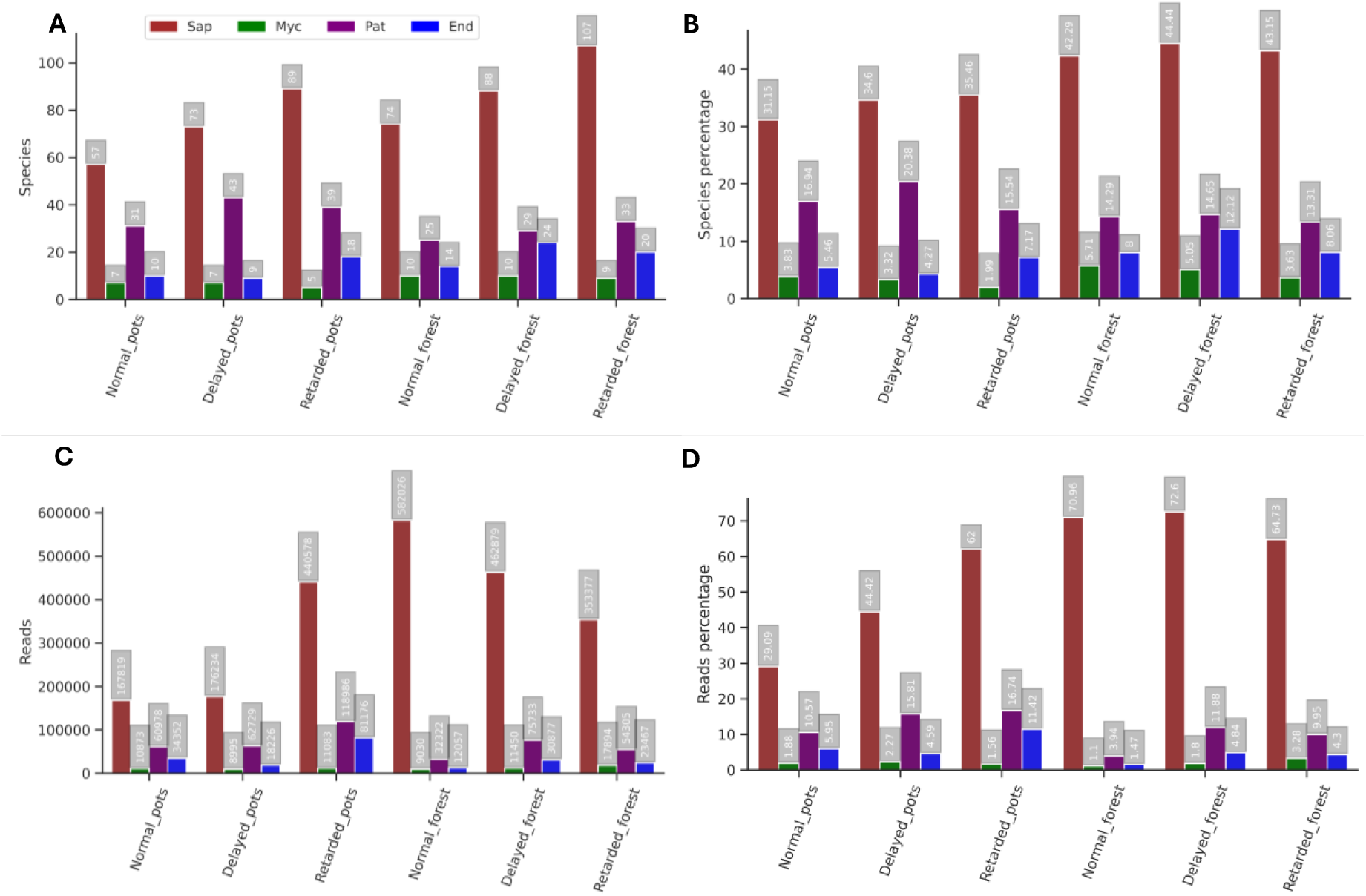
Ratios of plant related guilds in each sample set showing contribution in number of species (A), percentage of all species in sample set (B), abundance values (C) and abundance percentage of all reads in sample set (D). Values for species ecologically not assigned in FungalTraits database (ENA) vs all other guilds is shown on Figure 11.

#### 2.1.3. Stage III – Co-occurrence networks

Usage of co-occurrence networks have two major strengths. First is a unique approach toward analysis of species-species interactions both on level of singular relations as well as groups. Second is access to graph theory methods. That field of mathematics has wide range of formulas that can be applied to ecological data, exposing information that might be hard to access with other analytical approaches. An example network can be seen on a Figure 4 panel A.

**Figure 4.**
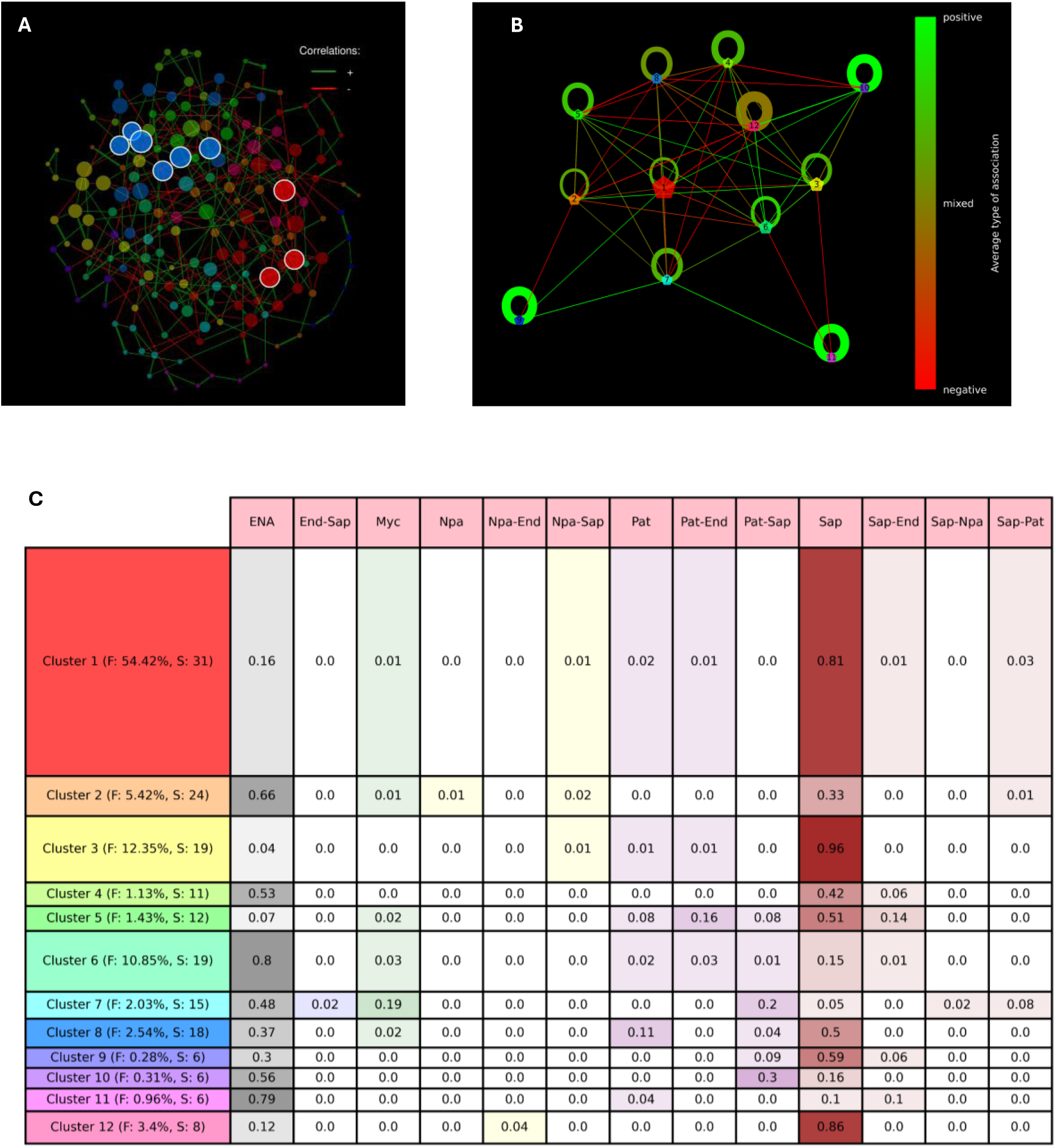
Co-occurrence network visualization of the Normal Forest, shown from three complementary perspectives: a species-level network generated using NetCoMi (A), a cluster-level network produced via ClusterCollapse (B), and a Make-up Table summarizing cluster trophic composition (C). Node colours in the species-level network (A) indicate cluster assignment and are consistently applied to cluster nodes in (B) and row labels in (C). Node size in (A) reflects a normalized centrality metric (eigenvector centrality), with hub nodes marked by a white border. In (B), node size corresponds to cluster abundance (min-max normalized). Edge width represents either pairwise association strength (A) or the density of collapsed edges (B). In the Make-up Table (C), cluster abundance is represented by row thickness, and row labels display total abundance (F) and number of species (S) per cluster. The trophic composition of each cluster in the Make-up Table (C) is represented by a two-part guild assignment, shown as percentages and emphasized through cell colours.

##### Network construction

The correlation of species occurrence in these graphs is calculated from the abundance patterns for each pair of species, which are only compositional, not quantitative due to nature of metagenomics. That demands use of tools that are prepared to handle such type of data. They must be aware of sparsity of metagenomic data. Thus, it is not proper to use direct correlation measures such as Pearson correlation coefficient or Spearman’s rank correlation coefficient without data transformation. For network construction, we chose NetCoMi R package (Peschel et al. 2021). It has a simple user interface and can pipe the input data into multiple different methods of network construction, from most basic correlation measures to ones specially created to handle metagenomic data. It performs pre-processing of data in form of zero-treatment and normalisation, followed by estimation of species-species associations, sparcification of the network, transformation into dissimilarity and then similarity which is the final edge weight. Two of association estimation methods: SPRING (Yoon et al. 2019) and SpiecEasi (Kurtz et al. 2015) do not require zero-treatment, normalisation or any data pre-processing as they were created specifically to handle microbial abundance data and perform those operations internally. We have decided to use SpiecEasi in GiaC. A key thing to note here is transformation of associations into dissimilarity. NetCoMi provides two methods following Dongen and Enright (2012): signed and unsigned. As final edge weight is 1 – dissimilarity, signed method gives the negatively associated taxa small edge weight and unsigned will give the strongly connected species high weight regardless of the sign. We have decided to use signed method.

##### Topology characterisation

Topology of a network simply means how nodes are connected by edges in a graph. That can be split into 3 parts: grouping species into clusters, finding nodes with special properties and edges with special properties. NetCoMi (Peschel et al. 2021) performs two of those tasks: cluster assignment and selection of unique nodes. For the search of special edges, we chose NetworkX Python package (Hagberg et al. 2008). Graphs are stored in object similar to native dict object in Python, which allows for very simple access to specific node or edge. This adds to multitude of graph analysis methods available already in the package, by allowing easy prototyping of custom methods, which we utilised in development of GiaC.

Clusters are found by one of several methods, such as modularity maximalisation method (Clauset et al. 2004) or hierarchical clustering. For GiaC method we chose modularity maximalisation. Modularity describes how much a cluster is a separate module inside of the whole graph taking into account not only binary existence of an edge between two species, but also the strength of that connection (Newman et al. 2003). With that approach, clusters represent groups of species that are much more connected with each other than with other clusters in the network. Unique nodes are labelled as hubs by NetCoMi (Peschel et al. 2021). Here we will refer to hubs only as nodes that have the highest eigenvector centrality (Bonacich 1972). This centrality measure states that the more central the node is, the more central are nodes connected to it. It does not account for abundance of a species that a node represents, meaning that even rare species can be flagged as special.

A most common type of a unique edge is a bridge. It is an edge whose removal will lead to existence of more parts of a graph with no connection to each other. A similar case exists for a node, here called a bridge node. It has the same property as a bridge. Bridges and bridge nodes are most basic way to measure how resilient to damage a network is. There is a caveat to this approach. A bridge can be near the middle of the network or in the periphery. A destroyed bridge near the middle will result in two big groups of disconnected nodes, while a destroyed bridge in the periphery will have lesser ramifications. At the end of this step, the resulting graph represents species as nodes, their associations (after all post-processing) as edges. Nodes hold information on how many reads were assigned to that species and what is the cluster and trophic guild of the species. Edges have their strength (the value of association) and sign as a colour (negative = red, positive = green) assigned. This graph has no selfloops, no multiedges and all edges are undirected.

##### Network evaluation

The final network can be evaluated in multiple ways. For example, existence of bridges or bridge nodes, what nodes were marked as hub or number of clusters. Except those options, there is a myriad of numerical parameters that describe specific characteristics of a given network. NetCoMi (Peschel et al. 2021) provides a list of parameters, here we described ones we deem most useful:

- Component number – number of separate intraconnected node groups in a graph
- Clustering Coefficient – global clustering coefficient, average of how densely connected is neighbourhood of a node, measures how clustered is the network, calculated according to Barrat at al. (2004)
- Modularity – already described above, worth noting is that weak positive connections inside a cluster will lower the value, measures how well the clusters are separated from each other
- Density – number of edges as percentage of all possible edges, given by Formula X, measures how connected a network is

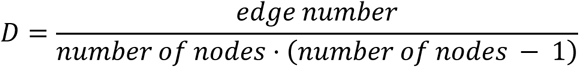
- Positive edge percentage
- Natural connectivity – average eigenvalue of adjacency matrix representing the graph, represents a weighted average number of closed loops of different lengths, measures how resilient the network is, especially to random edge removal (Wu et al. 2010)
- Number of clusters
- Vertex connectivity – minimal number of nodes that needs to be removed to disconnect network into two components. If there are more than 1 component in a graph, this will be 0
- Edge connectivity – similar to vertex connectivity, but for number of edges

Except those, we have selected several more useful parameters available in NetworkX:

- Number of isolates – number of nodes with no neighbours
- Diameter – number of edges between two most distant nodes in a network along the shortest path between those two nodes
- Radius – distance between the centre (node that is closest to every other node) and node furthest away from the centre

##### Graph coarsening – ClusterCollapse method

Clusters of nodes can be analysed by examining their external and internal edges as well as composition of the nodes within each cluster. For analysis of graph topology on the level of clusters instead of singular nodes we have utilised and fine-tuned a special type of graph coarsening based on edge contractions. We called this transformation the ClusterCollapse method.

Graph coarsening is a wide range of methods used to simplify graphs into smaller ones that still keep wanted information. Edge contraction is a process in which two nodes connected by an edge are transformed into a single node with all edges those two combined had, except the contracted one. Usually, this process does not represent contracted edges in any way, which can lead to loss of information if the process is repeated. We use ClusterCollapse method based on iterative edge contractions to collapse all nodes in each cluster into a singular node, keeping the contracted edges. For that purpose, we have chosen to retain the contracted edges inside a cluster as a self-loop. This way some information is retained, which we focused on: average strength, average type of connection (positive or negative in the original graph) and density of that connection.

Average of strength of a collapsed edge is a simple arithmetic mean. Density is calculated same as in “Formula X–1” where the value *number of nodes* is the node population in a given cluster for a self-loop or a population of the smaller cluster for an edge between clusters. We encode the average type of connection as a colour, utilising previously encoding positive and negative association as two different RGB colours. We represent each base colour in a RGB triplet as a decimal in range from 0 to 1. Encoding plus (green) as (0, 1, 0) and minus (red) as (1, 0, 0) gives us wide range of values. The colour is calculated as a mean of all colours that make this edge.

During normal edge contraction transforming one side of a triangle of nodes will produce a multiedge, as the new node has two connections to the same place. We decided to average multiedges in the same way as we average self-loop by producing a single edge between two cluster-nodes. Cluster-nodes (nodes representing whole clusters) hold number of reads assigned to all species in that cluster.

ClusterCollapse graphs simplify over 200 species into up to 15 nodes, which makes them easy to visualise. For that task, we have devised a template that shows cluster nodes as colourful pentagon with size (number of reads in a cluster) being normalised using min-max method. Thickness of an edge represents the density of the connection while the colour of a given edge is an average colour of a collapsed edge. See example on Fig 4 panel B.

It also has an added value of being by-the-way homogeneity test, what is described in detail in section 3.1.3.

##### Make-up table

Second half of cluster analysis is a visualisation called Make-upTable. It is a table-based visualisation representing a trophic composition of all clusters in a network. Example is shown on Figure 4 panel C. Make-upTable shows each cluster as a row with columns representing different trophic guilds. Here, we use full two-part guild acronym with primary lifestyle being the first half. Colours of guilds depend on the primary lifestyle assigned. First column of the table is the number of the cluster with their colours synchronised with ClusterCollapse and original graph visualisation made in NetCoMi. It also shows percentage of all reads that are assigned to each cluster, which is represented as thickness of each row.

### 2.2. Plant morphometry

The physiological state of tree seedlings is a dynamic interplay of genetic, environmental, and microbial factors, with the soil microbiome serving as a critical determinant of plant health and development (Furbank & Tester, 2011). State-of-the-art plant phenomics increasingly relies on remote sensing-based morphometric methods to assess growth and structural traits non-invasively at multiple scales (Abebe et al., 2023; Humplík et al., 2015).

Here we present the morphometric side of our methodology, optimized for experiments with limited resources. This method leverages remotely sensed tree crown traits, such as: green pigment content, size (area, perimeter, width, height), shape (compactness, roundness, eccentricity, isotropy), the parameters specific for used software (MorphoAnalyzer, Photon Systems Instruments, Czech Republic) and general development traits: number of branches, length of branches and number of leaves. This approach enabled to classify seedlings into phenotypic groups without relying on expensive molecular or imaging technologies (Araus et al., 2018). See Table 1 for mathematical definitions of parameters used in MorphoAnalyzer software for evaluation of top or side tree crown images.

**Table 1.** Morphometric parameters calculated using MorphoAnalyzer software selected for present analysis and their mathematical definitions.

| Parameter | Definition | Formula |
| --- | --- | --- |
| AREA_PX | Total area covered by the plant | Count pixels |
| PERIMETER_PX | Length of the plant perimeter. | Count pixels |
| ROUNDNESS | Ratio between area and perimeter of the plant surface | $\{4 * \pi * \text{Area}\} / \{\text{Perimeter}\}^2$ |
| COMPACTNESS | Ratio between area and surface of the convex hull enveloping the plant. More "compact" plants have shorter petioles and massive leaf blades | $\{\text{Area}\} \text{ over } \{\text{ConvexHullArea}\}$ |
| HEIGHT_PX | Height of the bounding box enveloping the plant | Count pixels |
| WIDTH_PX | Width of the side bounding box enveloping the plant | Count pixels |
| ISOTROPY | Isotropy of the object, computed as ratio of area and perimeter of the polygon created by object vertices | $4 \times \pi \times (\text{Polygon\_Area}) / (\text{Polygon\_Perimeter})^2$ |

#### 2.2.1 Image Acquisition preprocessing and Transfer

In presented methodology, morphometric analysis of the plants was performed with use of imaging based on the RGB colour model (Sadeghi-Tehran et al., 2017). Our test images were captured using a smartphone (Huawei P40 Lite). In order to standardize image taking procedure all pictures were taken at fixed distance between camera and plant, fixed focal length and same resolution settings (Lee et al., 2018). Capture of top-view and side-view images under natural lighting conditions was performed in similar time frame for consistency. A key point in image acquisition is the removal of background noise (Okyere et al., 2023). In our approach we used black agrotextile sheets to minimize this effect, but light refractions could not be completely avoided. We believe that usage of material with negligible reflective potential or use controlled lighting (e.g., lightbox) for future imaging would eliminate this hindrance. Due to high morphometric variation not only between groups but also within, we supplemented remote sensing with manual measurements. In order to ensure reduced number of variables affecting image analysis, use of smartphone camera filters is not recommended also in presented methodology we did not perform post-processing of images. We did however preformed JPEG to PNG format conversion to match image analysis software required input format.

#### 2.2.2 Mask construction and colour segmentation

We preformed image analysis using MorphoAnalyzer software (see file Supplementary_3_morphometry_data). Process begins in Mask Builder module, here user can define region (binary mask) in picture where software will perform further operations (Henke et al., 2021). Following it, original image and primary mask can be forwarded to Plant Mask Tool module. This module allows to create regions of interest (ROIs) within space defined by the primary mask. One does so by choosing colours that correspond to areas of interest (wanted colours) and the background (unwanted colours) (Li et al., 2021). Based upon them a Colour Formula is calculated, using it software forms a secondary mask that cuts object of interest within space defined by primary mask from background. Colour formula coupled with other parameters affecting secondary mask such as threshold, reflectance reduction, median filter size, min object size, min hole size allows software for precise cut out virtual silhouettes of object of interest, such as analysed plant, from leftover background (Lee et al., 2018).

Next step of image analysis is a colour segmentation and selection of output parameters. For it software uses defined ROI and its own set of colour definitions. However, it is imperative to optimize colour definitions towards conducted experiment. Definitions are stored as a TXT file and can be download, edited and uploaded as such. Following defining colours, selection of analysed parameters is required. These parameters can be separated into two distinct categories Top View and Side View with each of them having their own set. Using MorphoAnalyzer software, we have access to side view parameters such as Area, Perimeter, Compactness, Width and Height. Top View parameters include Area, Perimeter, Roundness, Roundness 2, Isotropy, Compactness, Eccentricity, RMS and SOL.

#### 2.2.3 Image analysis

With parameters chosen and colours defined software offers two modes for preforming image analysis: single run and batch run. Batch run allows user to preform mass analysis of whole sets of images here called batches. After choosing input folder with images properly tagged, it will analyse each image using the same defined space but self-building ROI. This mode will optimise construction of plant mask based on colours of interest requiring it to have highly standardized sets of images. The image tags will also assign image to the numerical results in output table. The other mode, single run mode performs analogous operations with a key difference that each image is analysed separately allowing user to assist and monitor software ROI selection. Other crucial difference in single run mode lies in its option to preview the original image, plant mask and colour segmentation output. Single run mode also requires user to manually feed it images and masks for analysis.

## 3. Methodology effectiveness

In this section, we present first detailed explanation and justification of GiaC method, then in second subchapter same for plant morphometry. We outlined results of our holobiont assessment in section 3.3. In that part we also show if the GiaC+phenomic approach was able to test hypotheses outlined before.

### 3.1. GiaC

#### 3.1.1. Stage I – Taxonomic and trophic assignment

As it is already stated in section 2, quality of results for taxonomic identification, functional assignment and filtering depends on methods, tools and references available. Here we list several general ideas of what types of methods would be useful for what task. We also describe the effectiveness of F99 filtering and usefulness of Sankey diagrams (Kennedy and Sankey 1898). All specific values and details of results for our test dataset are presented as Sankey diagrams on Figure 1 (see Supplementary_1_sankey_flows file for all charts not included in the main text).

##### Metabarcoding, quality control and barcode sequence grouping

For this proof-of-concept study, we have used DNA using DNeasy PowerSoil Pro Kit (Qiagen 47014) for isolation and Ion GeneStudio S5 System (Thermo Fisher Scientific) with KYO-ITS primers (Toju et al. 2012). We have utilised Qiime2 quality-filter plugin with flag set to p-min-quality = 10 for QC and DADA2 plugin for ASV creation with parameters set to p-min-quality = 0 (Bolyen et al. 2019; Callahan et al. 2016). This generated 7 239 ASVs in the whole dataset. Using long reads (Nanopore, PacBio) or multiple barcodes could also be used with our methodology, as that does not change later stages of GiaC, as long as QC and ASV tool are adjusted to that data.

##### Taxonomic and functional assignment

At the time of writing this paper most precise and still properly moderated database for taxonomic identification is UNITE (Kõljalg et al. 2005; Kõljalg et al. 2019). It is the go-to database for most cases, and its continued development only adds to the value. Even though there are many different complex tools that can be used to identify species by their barcode, BLAST (Altschul 2005) can still be used. There are even multiple tools that implement BLAST-like algorithms for taxonomic identification. For our dataset we have used UNITE as reference database and BLAST for taxonomic identification. That assigned 6656 ASV to fungal species in one or more samples.

In this proof-of-concept study, we have checked the overlap of FungalTraits and FUNGuild databases for functional assignment. At the time of writing, they both agree on functional roles for 53.86% species, partially agree on 22.89% species and differ only in 23.25% species for test dataset of 1372 species. Taking this example into consideration and what was already reported (Tanunchai et al. 2023), we advise using FungalTraits. A further source of data for functional identification may be a search for information on whether species genomes contain genetic markers which are specific to certain ecological role. This however requires existence of those genomes in a database and knowledge of what genetic markers are specific to what ecological role.

##### Species filtering - Flexible99

F99 method is not much more complex than singleton removal. Singleton removal focuses only on species or ASVs with single occurrence. That fixed cut-off value can be increased to 3, 5 or even 15 occurrences to remove ASVs and taxons that may be a contamination. While we use a flexible cut-off that is automatically adjusted to a given dataset, in all cases except extreme ones, it will be much higher than 1. Thus, our method not only removes those unsure species, possible contaminants, but also focuses downstream analysis on more abundant species. This removes only 1% of all reads in a dataset but results in removal of around 40% of species (Fig. 1). It can be argued that it loses information on rare species, but as the focus of this methodology is to assess general trends in fungal communities, it is very unlikely that very rare species with abundance on the level of contamination will have any visible impact.

##### Sankey diagram

The Sankey flow diagram template we created manages to keep the overview simple yet informative. As one can see the Fig. 2 is informative enough to exclude any numbers or statistics from description of Stage I except the number of ASVs created. We strongly advise to use this diagram with our template for clear, concise report on data filtering in metagenomics. We also advocate wide adaptation of Sankey diagrams in all data analysis in biology. In this study we utilised SankeyFlow Python package (Xu, 2022).

#### 3.1.2. Stage II – Microbiome composition analysis

The visualisations we have listed in section 2, are not without their limitations. Users of those tools or whole GiaC method have to be aware of several ways in which they can be misleading or hard to interpret.

##### Clustermap

Clustermap being a combination of dendrogram and a heatmap is a more comprehensive view than other shown in this Stage. Using percentile normalisation amplifies existence vs absence of a species but does not represent absolute values. Clustermap is a method with less limitations than other steps of Stage II but its complexity makes it harder to interpret.

As previously mentioned in section 2.1.1, clustermaps emphasises internal difference in a sample set, not compare separate sets. That requires treating multiple sample sets as a singular combined one. We advise to treat them as one set already on F99 filtering stage. This method adds up occurrences from all samples for each species. That results in keeping more species with low abundance or found in very few samples than if two combined sample sets were filtered separately, giving some insight into occurrence pattern of rare species. F99 also keeps species unique to one part of the combined sample set. Nevertheless, it will still remove species with very low abundance in both parts of the combined sample set.

Biggest pitfall of using a clustermap is reading the dendrograms. The method show patterns in occurrence through reordering of rows and columns, putting similar ones next to each other. Dendrograms are products of hierarchical clustering methods, which work with assumption that each cluster should be split into two subgroups. That will lead to a confusing result in cases where that assumption is wrong at some step. Dendrograms will exaggerate similarity between samples in cases of more than two subgroups.

Let’s take for example our dataset. We have two sites (Pots and Forest), each comprising three plant conditions. We can expect the top cluster to be partitioned into cluster of all Pots samples and second one of all Forest samples. This is well represented and can be seen on Figure 5. It however does not inform how similar the samples are. It might be a major difference or barely any. That feature of a clustermap has to be remembered, as it can be a source of confusing but valuable information. In our case clustermap did not partition each site by plant condition, what may be due to different species composition in sites being much more pronounced factor.

**Figure 5.**
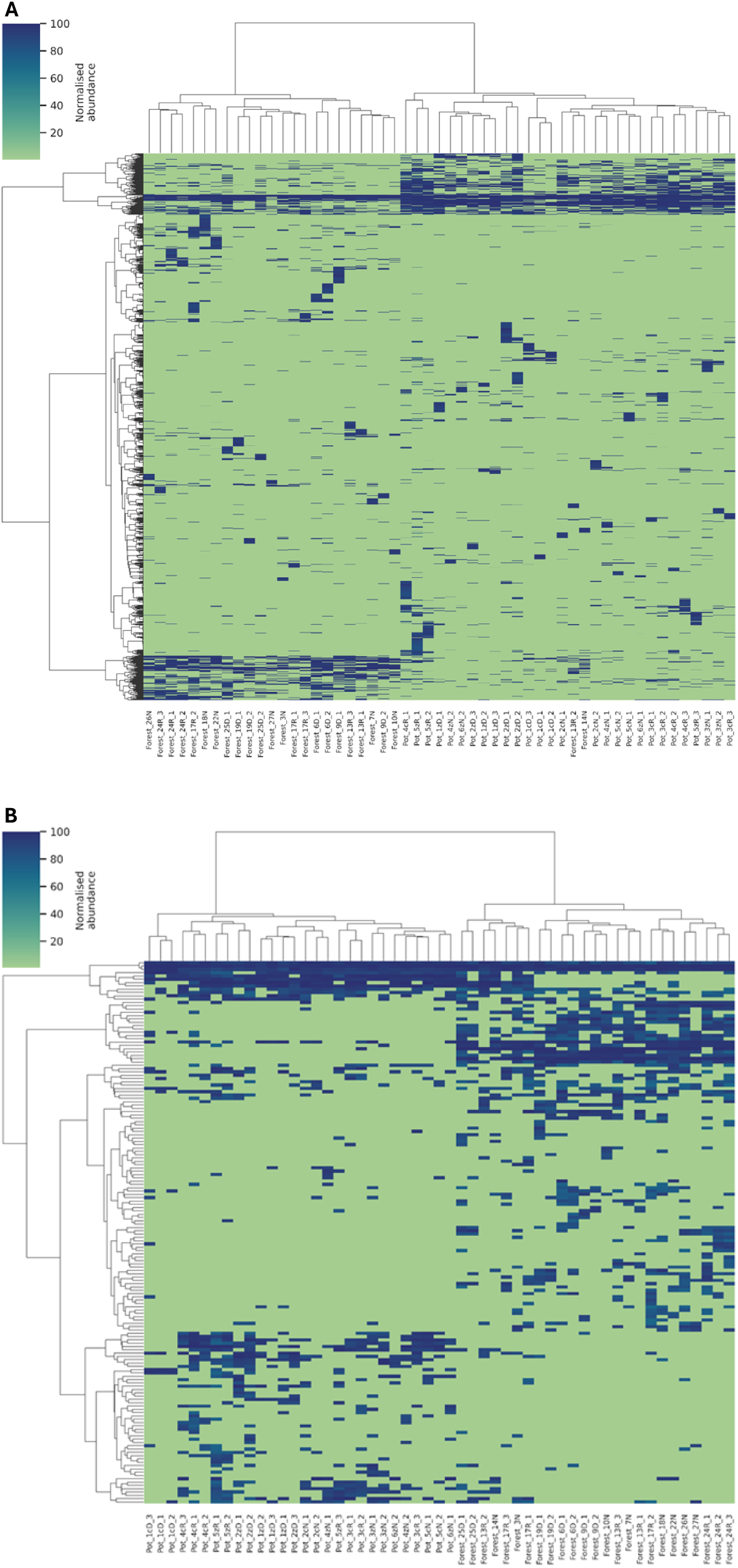
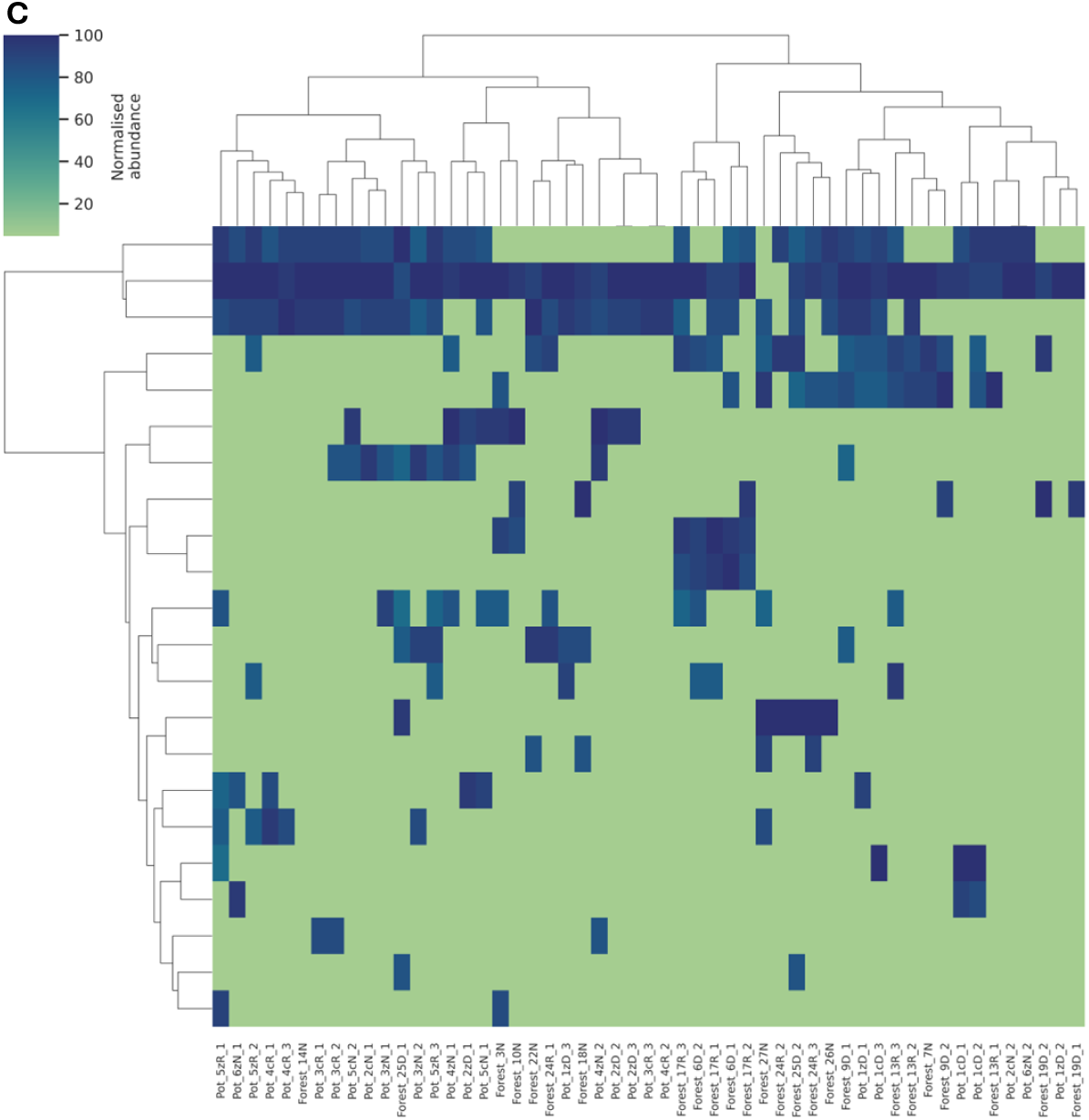
Clustermap visualisation of species abundance for each sample (colums) and species (rows) created using Ward’s minimum variance method and percentile normalization of abundance. Both clustermap of all species (A) and saprothropic species clustermap (B) separated samples into Pots and Forest. Mycorrhizal species (C) clustermap does not show a full separation.

#### 3.1.3. Stage III – Co-occurrence networks

The present section is devoted to clarifying certain intricate elements of Stage III of GiaC that were outlined in the main description of the methodology. We expand here on effectiveness of certain solutions or their biological interpretation.

##### Modularity

Using complex, state-of-the-art methods of microbiome co-occurrence network construction requires careful understanding of how they produce final graph. As we described before, after building association matrix we used signed formula for transformation into dissimilarity and modularity maximalisation approach for cluster assignment (Dongen and Enright 2012). This combines in a specific way with the way NetCoMi (Peschel et al. 2021) implemented calculating similarity and it tends to produce specific type of clusters. Because final edge weight is 1 – dissimilarity, signed method of transformation into dissimilarity will result in negative edges having lower weight than positive. Weak edges produce lower final value of modularity network parameter, as long as that edge have nodes on both ends in the same cluster (Newman et al. 2003). Strong green edges will increase modularity value more, as long as they are inside a cluster. Thus, using signed method results in a higher value of modularity if negative edges are between cluster. That characteristic can be leveraged together with modularity maximalisation as method of cluster assignment (Clauset et al. 2004). As the name describes, modularity maximalisation groups nodes into clusters with highest modularity. As modularity is higher when more edges with negative association are between clusters, modularity maximalisation will produce clusters that primarily have internal edges positive. Such clusters may represent groups of species cooperating in a studied site with negative interactions mostly placed between those groups. It can still keep some negative edges inside clusters, especially weak ones. This can be seen on a Figure 4 panel A.

##### Measuring network resilience

Network resilience can be measured in respect to two types of attacks: targeted or random. Targeted attack means that removal isn’t random, but of specific nodes or edges. Most sensitive to this attack are bridges and bridge nodes. NetCoMi (Peschel et al. 2021) implements natural connectivity parameter, which is not only a much better way to measure the resilience against targeted attacks, but also random ones (Wu et al. 2010). In our dataset the worse the plant condition, the lower the value of natural connectivity and number of bridges and bridge nodes was (see Supplementary_2_networks file). In contrast, better plant condition had networks with more bridge and bridge nodes and lower density but higher natural connectivity. Most of the bridges or bridge nodes were on the very periphery of the network with none connecting two clusters. This shows that a simple information of how many sensitive edges or nodes there are, does not represent how well a network is connected. It is crucial to know the position of a bridge. The more central the bridge is, the more chaos will removal of that edge result in. It is also important to remember, is that ecological networks are not static in time and change optimising to its environment. What might be a bridge in a given snapshot does not have to be a bridge in a different time point and removal of a bridge does not mean a network won’t reconnect easily. Non-static nature of ecological networks requires bridge analysis taking into account different time points, not one singular snapshot. All this taken into account requires use of multiple resilience measures concurrently when analysing co-occurrence networks, which is part of GiaC method.

##### Encoding average association type in ClusterCollapse

Type of a species association (single edge) in a co-occurrence network can be positive (+) or negative (−). It is a discrete, binary variable, which limits values to only two possible with nothing in between. Thus, there is no straight-forward way to represent an average type when trying to combine multiple edges. A good solution is to encode the type as a variable with range, giving us an option to average like any other continuous variable. We decided to encode the type as a colour in RGB model for two reasons. Firstly, colours are triplets of number with multiple values possible, which gives us a range. Secondly, colours can also be used for a visualisation on a graph drawing (Fig 4 panel B) and are still simple numbers, easy to analyse in code. We picked decimal representation in range 0-1 as it is a continuous variable, not discrete like integer 0 to 255. If we used 0-255 encoding, this would allow only 256 options, but using 0-1 range gives us magnitudes more. This encoding gives ClusterCollapse ability to keep as much information as possible from all contracted edges.

##### ClusterCollapse as a homogeneity test

ClusterCollapse has an added value of being a by-the-way homogeneity test. What is counter intuitive a more regular network means more internal differences in analysed sample set. Figure 6 panel A shows a ClusterCollapse network made from all samples in our study. The network is a completely connected graph, made up of almost all connections fully green (88.39 positive edge percentage for the original SpiecEasi graph) and has only 5 cluster-nodes. This is very different from a network modelling microbiome of Normal plant condition in Forest site, which is shown on panel B of Figure 4. That one has more cluster-nodes (12), more edges of mixed or red type (61.13 positive edge percentage for the original graph) and many cluster-nodes are not directly connected with the rest of the network. Panel B of Figure 6 shows a graph of Normal condition from both sites mixed together. It’s visible that this network has characteristics midway between network of all samples and Normal condition in forest site. It has 10 cluster-nodes, most but not all edges are fully green (81.26 positive edge percentage in the original graph), but all cluster-nodes are directly connected. This shows that ClusterCollapse network transformation is able to magnify graph characteristics that change with the level of homogeneity in the sample set. This is especially useful trait of the method, as ClusterCollapse primary role is not to test for homogeneity. That trait allows it to perform a dual role at the same time, simplifying whole analytical process.

**Figure 6.**
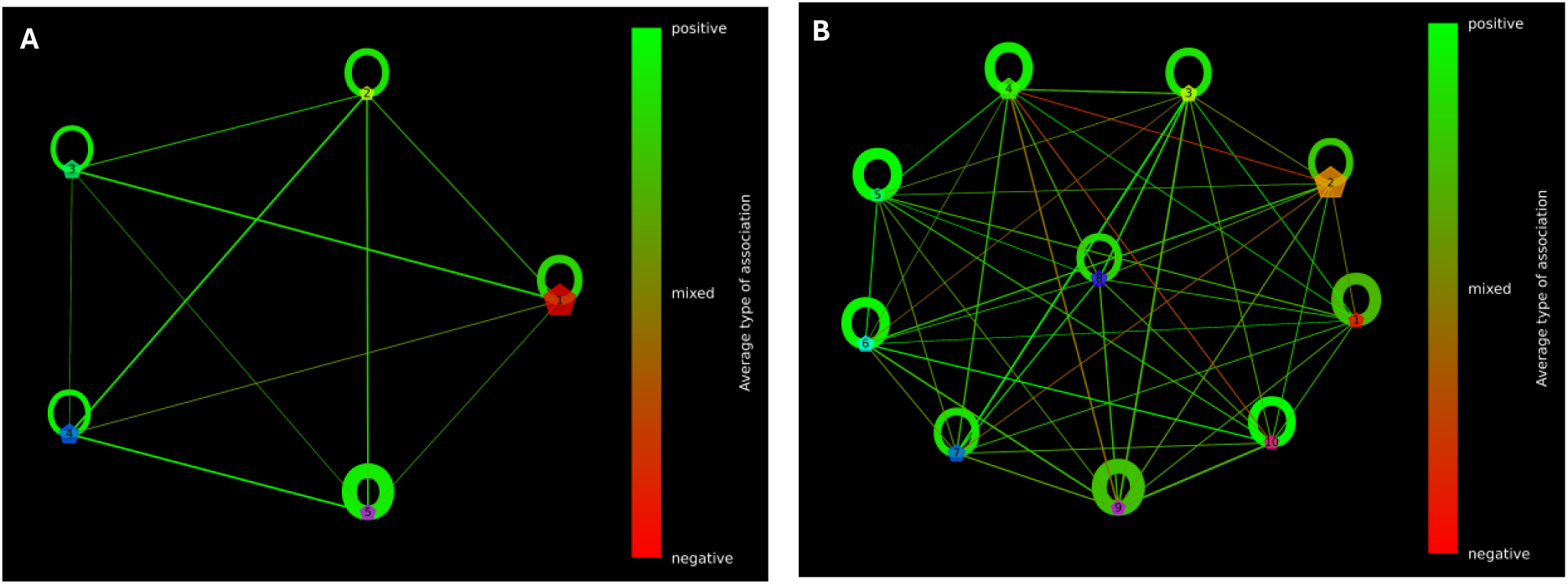
Demonstration of how ClusterCollapse can be used as homogeneity test. Here shown are two different, non-homogeneous sample set: all samples (A) and Normal plant condition from both sites combined (B). For comparison, a homogeneous set can be seen on Fig 4 (panel B). For details on how the test can be performed, see section 3.1.3. under “ClusterCollapse as a homogeneity test”. For guidance on interpreting the graphs, see Fig. 4.

### 3.2. Plant morphometry

Most crucial issue of our morphometric analysis has proven to be light conditions of image taken (Sadeghi-Tehran et al., 2017). While capturing images of trees under natural field conditions, variations in lighting, its refractions, leaf angles, and background complexity introduced challenges that required careful countermeasures in image taking and data processing (Okyere et al., 2023; Henke et al., 2020; Xia et al., 2015). We succeeded in reducing background noise influence and proposed additional steps that will allow optimisation of future experiments. Light refractions and reflections had proven to be most difficult issue for successful image analysis in field conditions (Henke et al., 2020; Sadeghi-Tehran et al., 2017; Xia et al., 2015). Remaining issue were leftover reflections on background material caused by wind induced movement differing from picture to picture. We resolved this issue using Single run mode. Single run approach while more time consuming has proven to be a preferable method thanks to its options allowing parameters adjustment for every plant analysed. We suggest utilization of dedicated portable/mobile non-reflective background preferably made of heavy or solid material such as light box. Utilization of such device would reduce refractions and reflections issue to point of irrelevance.

#### Colour Segmentation Limitations and Mitigation Strategies

Main method of image analysis in presented work is RGB colour scheme-based colour segmentation (Humplík et al., 2015; Sadeghi-Tehran et al., 2017). Colour segmentation, while a powerful tool, does faces several limitations that limits its applicability in presented field experiment approach (Okyere et al., 2023; Sadeghi-Tehran et al., 2017). One major challenge is the inherent sensitivity of colour-based segmentation to variations in lighting conditions (Chopin et al., 2018; Henke et al., 2020; Okyere et al., 2023). Software’s operating with this method do not differ colours same as user and image taking device with RGB, they generally define them as a part of mathematical formula. Because of this, variation in lighting conditions during subsequent measurements may lead to inconsistent results in the reported set of hues (Sadeghi-Tehran et al., 2017; Yu et al., 2017). Additionally, colour similarity between objects of interest or transitions between them and background elements can cause misclassification, where distinct regions are erroneously grouped together (Henke et al., 2020). Shadows, reflections, and subtle gradations in colour further complicate segmentation by introducing noise or artificial boundaries (Henke et al., 2020; Sadeghi-Tehran et al., 2017).

There are several strategies can be employed to mitigate these issues such as colour normalization, which while it does enhance ROIs borders definitions it introduces additional colour distortions making it not suitable in all scenarios, including those presented here (Cai et al., 2016; Chopin et al., 2018). Other methods also proven to be varied but much more stable in their results. Standardising illumination improves results, but applying it in field conditions is complicated without dedicated equipment. (Sadeghi-Tehran et al., 2017; Yu et al., 2017). Applying advanced colour spaces or reflectance reduction requires an understanding of image analysis software algorithms, otherwise negative results may be obtained (Altukhov, 2022; Henke et al., 2021; Okyere et al., 2023). Utilizing machine learning-based approaches on other hand requires significant datasets for training making it of limited value for presented experiment (Henke et al., 2020; Okyere et al., 2023).

Considering the above, in order to enhance consistency of the results by increasing contrast and reducing noise, we have employed combination of: minimum object/hole size definition, reflectance reluctance, and adaptive thresholding method offered by MorphoAnalyzer software. Despite the tool yielding somewhat varied results, likely attributable to light noise in the field, it prevented the analysis of less than 4% of the images captured.

### 3.3. Test data results

#### 3.3.1 GiaC

##### Species composition

Clustermap grouped samples together based on site they are from, not plant condition (Fig. 5). It shows that there is much bigger difference in species composition between sites than between conditions in the same site. This result is confirmed by Venn diagrams (Fig. 7) showing that each site shares a lot of species between all conditions, but the same condition in two sites shares only a small number of species. Venn diagrams also show increase of overall species number and condition specific species as plant condition worsens.

**Figure 7.**
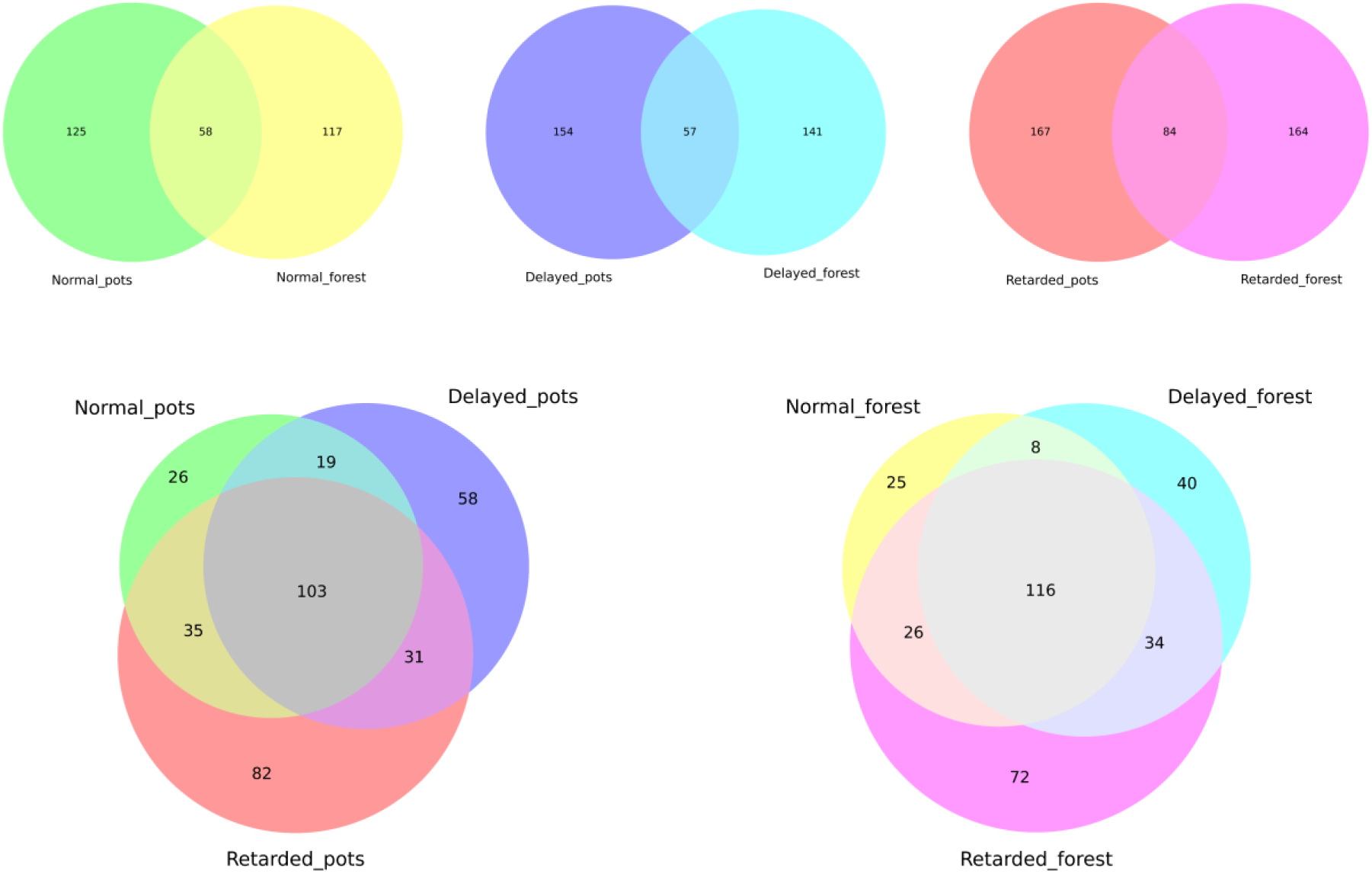
Venn diagrams showing numbers of specific and shared species between Pots or Forest sites for the same condition (top panels) or between different plants conditions (Normal, Delayed or Retarded) in the same site.

##### Trophic composition

Guild ratios show that the increase in number of species with worsening condition is mostly the increase of Sap guild (Fig. 3A) and ENA guild (Fig. 8A) in absolute numbers. Myc guild slowly drops, and other guilds show non-linear change. Percentage ratios show that the share of every guild except Myc and Sap in Pots fluctuates around the same value with no specific trend (Fig. 3B).

**Figure 8.**
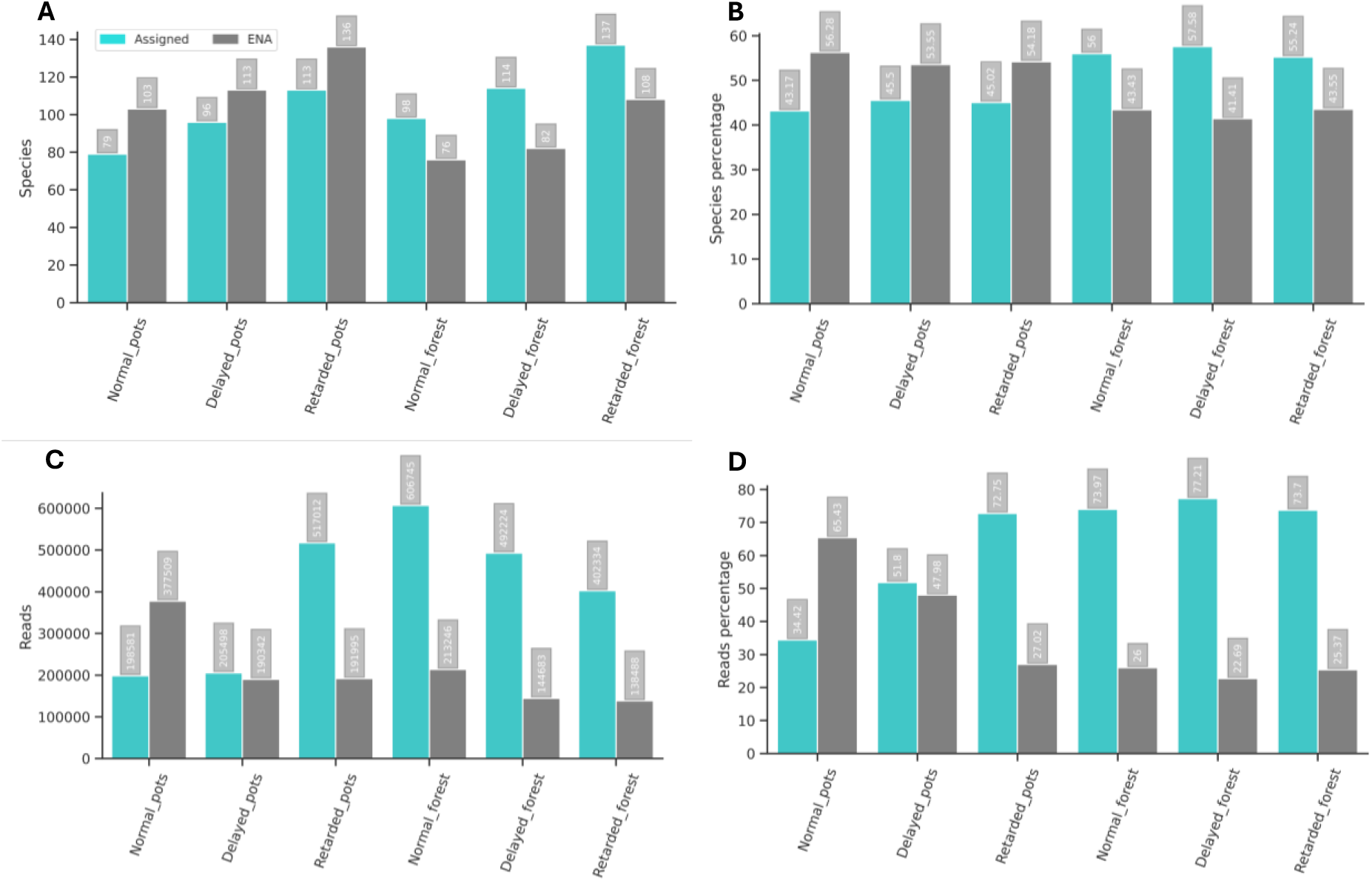
Ratios of plant related guilds in each sample set showing contribution in number of species (A), percentage of all species in sample set (B), abundance values (C) and abundance percentage of all reads in sample set (D). Values for species ecologically not assigned in FungalTraits database (ENA) vs all other guilds is shown on Figure 11.

The values of guild abundance display divergent trends with declining plant condition across both sites. Pots dataset is characterised by non-linear increase in reads of Sap and Pat guilds (Fig. 3CD) as well as non-linear drop of ENA reads (Fig. 8CD). Other guilds do not show any general trend. In Forest samples worse condition means linear drop of Sap, non-linear increase of Myc, Pat and End and non-linear drop of ENA species. ENA species warry in behaviour between site, in Forest always being around 25% of all reads, while in Pots dropping from 65.43% of reads in Normal samples to 47.98% in Delayed and 27.02% in Retarded. This shows that fungal species that support plant growth in isolated environment of pots are not well studied yet. Delayed and Normal conditions differ in terms of guild abundance between sites and more similar are Retarded sample sets from two sites.

##### Network evaluation

To visualise network parameters for 6 sample sets in our test data, we have used radar charts shown on Figure 9. Values on panels A and B are normalised among all 6 sets, but panels C and D are normalised inside a single site. This approach shows that while mycobiomes of plant of the same conditions from two sites differ in absolute values, the same condition from two sites share the tendency to have highest or lowest relative values. It allowed to correct for site specific characteristics and describe it not as having a certain absolute value, but having certain value being highest or lowest. There is a difference between conditions in the same site. There is also a big similarity between the same condition from different sites as seen on panels C and D. Fungal networks of Normal plants always have the sparsest graphs (highest values for radius, diameter, bridge nodes and bridges numbers) but most resilient to edge or node removal (highest natural connectivity) with most different clusters (highest modularity). Retarded networks tend to be densely connected (highest clustering coefficient and edge density) but have most similar clusters (lowest modularity) and are least resilient (lowest natural connectivity) despite less bridges. Delayed networks have characteristic of in-between state. The most similar are Normal condition networks, Retarded networks show a similar trend but Delayed behave a bit differently in each site.

**Figure 9.**
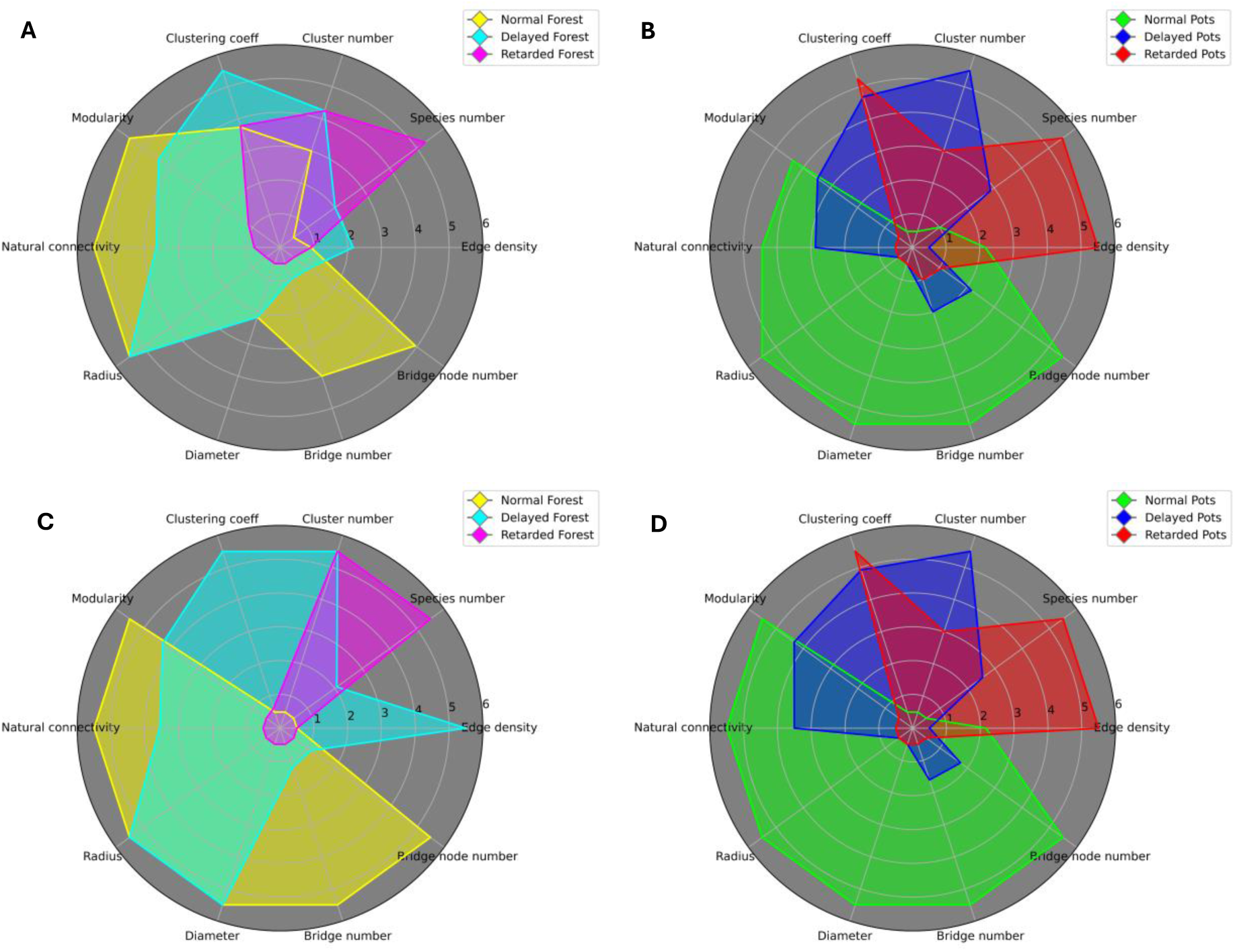
Diverging network parameters for Forest and Pots sample sets, normalized. Panels A and B show values normalized between all 6 sample sets, which allows for easy comparison. Panels C and D show the same values normalized in single site, separate for Forest and Pots.

##### Summary results

The results can be summarised to increase of chaos in mycobiome with worsening plant condition. In Forest dataset it is trend of decreasing summed abundance and increase in species number leading to microsites with many species of low abundance (Fig. 8). In Pots dataset there is no trend of abundance values, as different guilds show opposite behaviours, but worse condition has higher species number. Network parameters also show increase of chaos, as co-occurrence graphs in worse conditions are less structured. It is represented in lower modularity (less different clusters), bridge or bridge node number, radius and diameter as well as higher edge density in worse conditions. In Pots the Delayed samples more closely resemble the Retarded sample set (e.g. clustering coefficient), while in Forest Delayed aligns with the Normal condition (e.g. radius). It shows different behaviours of Delayed plant + mycobiome holobiont in each site.

#### 3.3.2 Morphometry

The calculated numerical plant parameters, such as crown size, shape, area and pigment content, number of leaves, branch length and foliage density, allowed the seedlings to be classified into three different vitality groups (Fig. 10, Fig. 11, Fig. 12). The most divergent indicators we aggregated with network parameters in the form of a summary radar chart (Fig. 13). For details of raw morphometric data see ‘Data Availability’ section.

**Figure 10.**
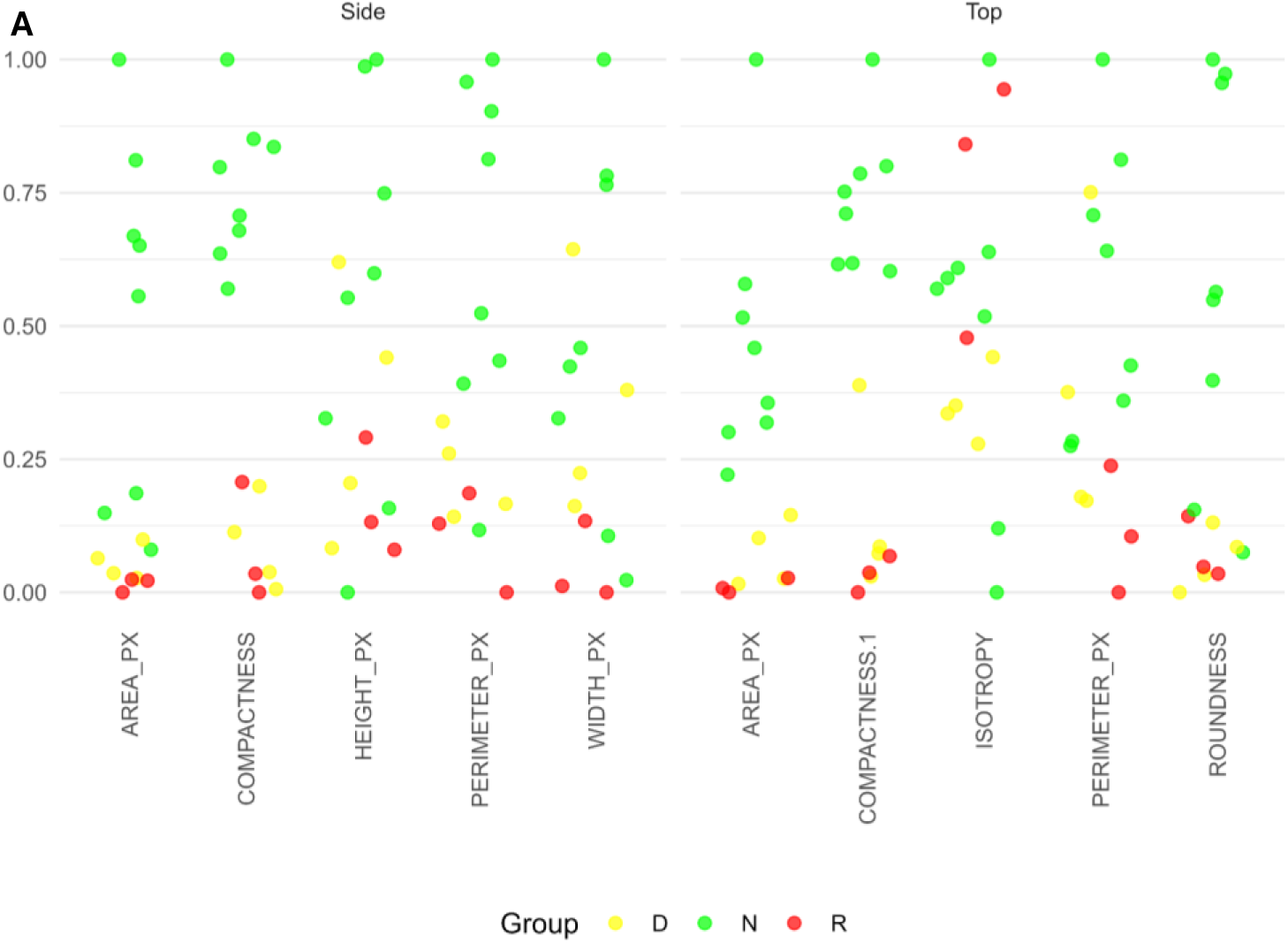
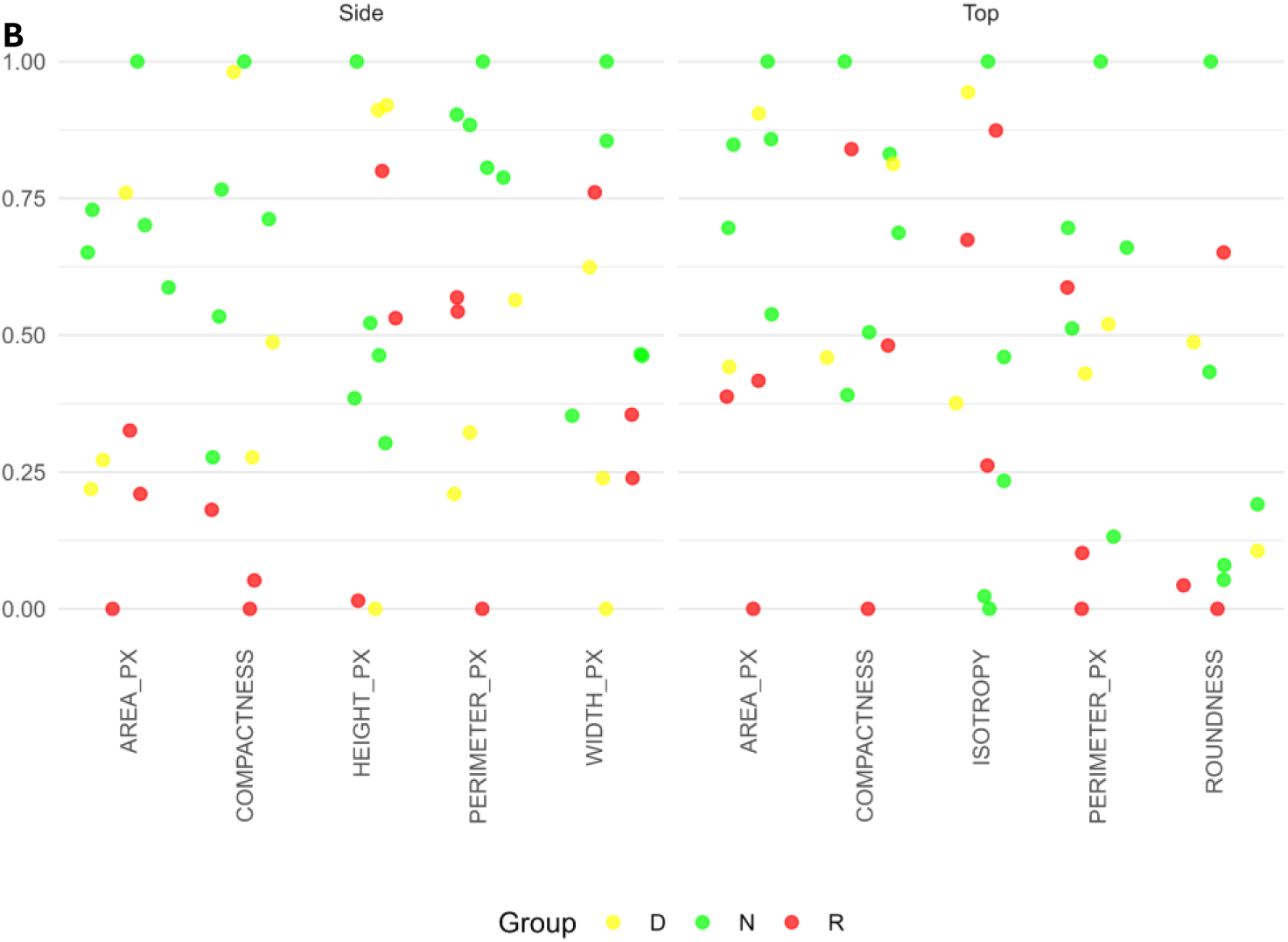
Morphometric analysis of RGB images (side and top views) of tree crowns used to classify seedling vitality in forest (A) and nursery (B) sites. The colours indicate groups of seedlings from specific site with clusters of parameters showing differences in growth delays: green-Normal (N), yellow-Delayed (D), red-Retarded (R). The variation in the computed shape and size descriptors (see formulas in Table 1) was normalised relative the Forest or Pots dataset and separately for side or top camera views.

**Figure 11.**
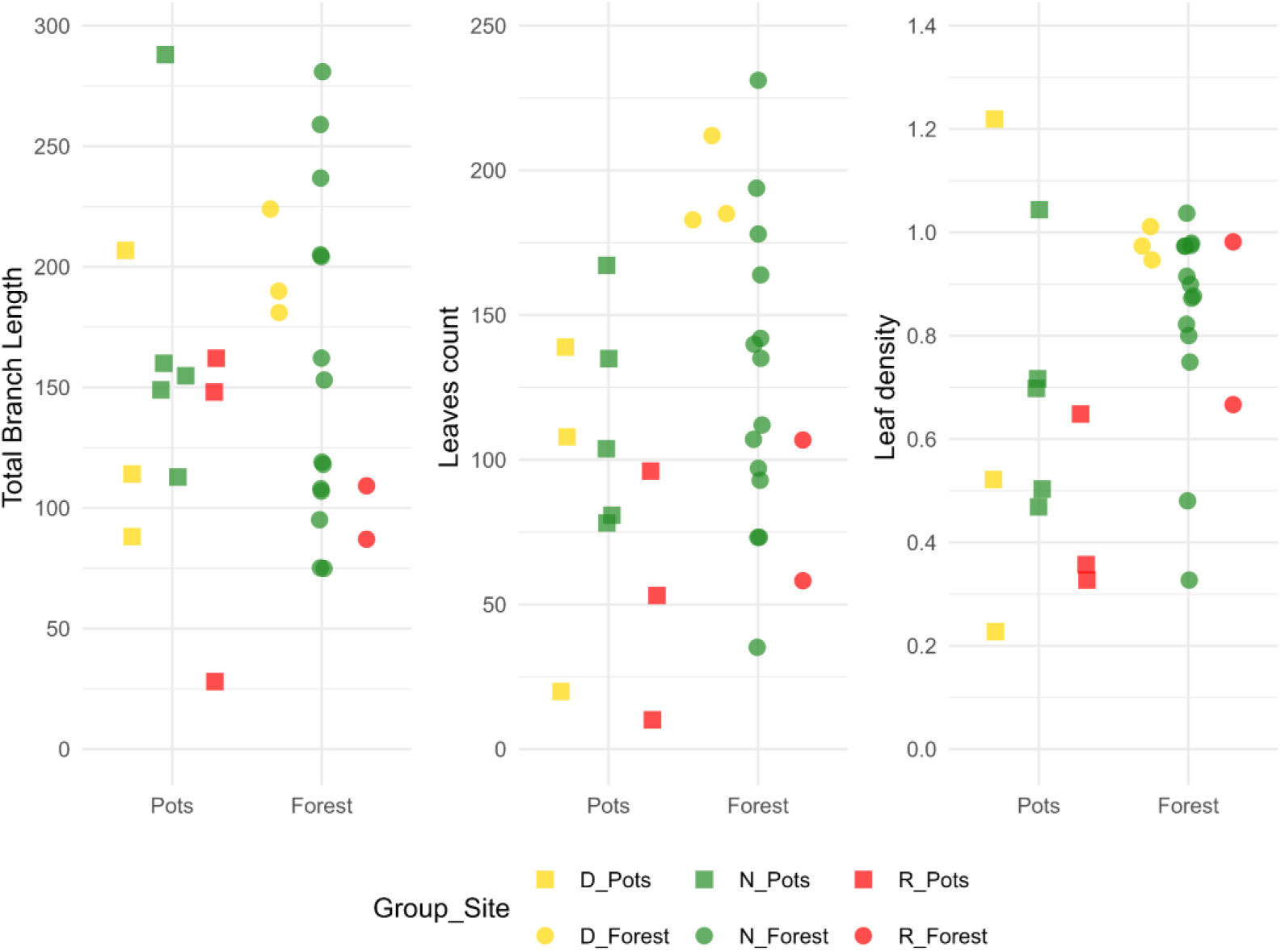
Manual metrics of plants from forest (circles) and nursery (squares) sites. The colours denote groups of seedlings from specific site with clusters of parameters indicating differences in growth delays: green-Normal (N), yellow-Delayed (D), red-Retarded (R). Values are plotted on the raw measurement scales (branch length in units reported in the dataset, leaf count as absolute counts, leaf density as the per-plant density metric), with each point representing one seedling observation.

**Figure 12.**
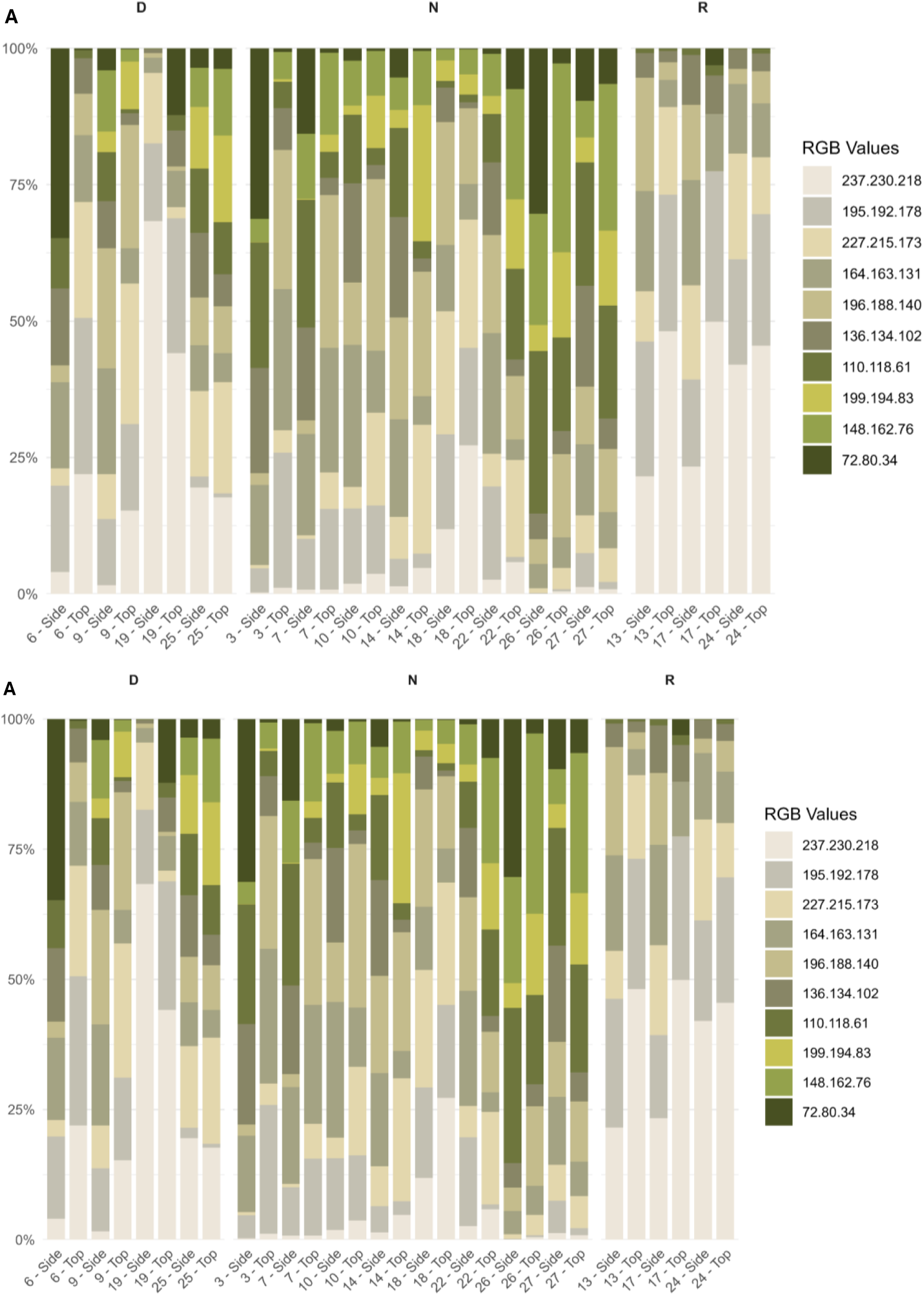
Classification of seedlings in forests (A) or nurseries (B) into groups differing in vitality according to variation in pigment content in the crown. RGB top and side tree images were analysed to calculate the area percentage of leaf pigments (green tones) and branches (gray colors) using colour segmentation and pixel count based methods. Proposed groups of different growth delays were indicated as: Normal (N), Delayed (D), Retarded (R).

**Figure 13.**
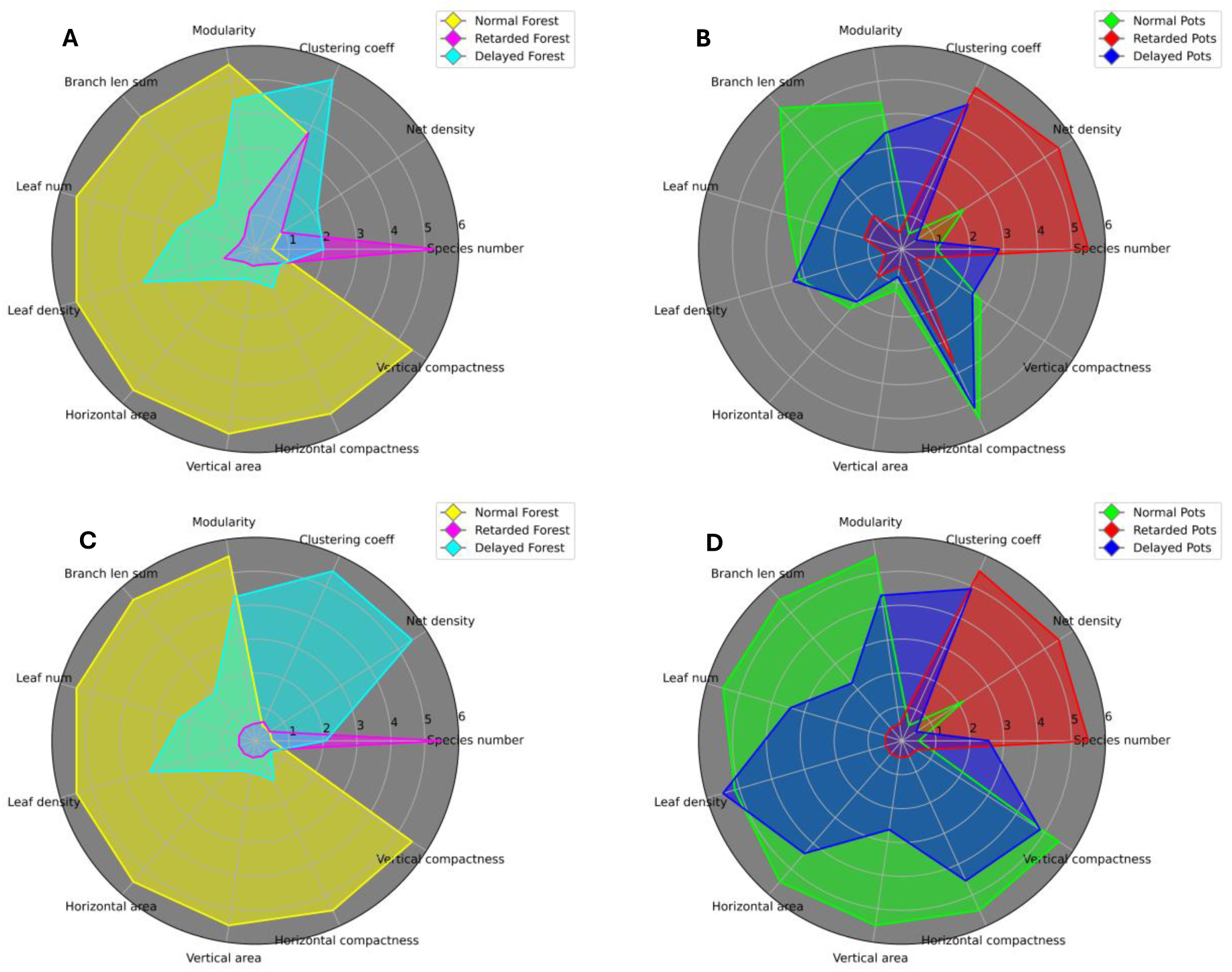
Diverging holobiont parameters (both morphometric and network) for Forest and Pots sample sets, normalized. Panels A and B show normalization between all 6 sample sets, which allows for easy comparison. Panels C and D show the same values normalized in single site, separate for Forest and Pots.

The “Normals” group (N), while exhibiting a full range of values across the tested parameters, consistently maintained at least half of its scores in the upper 50% for each measure. In contrast, the “Retarded” group (R) predominantly fell within the lowest 25% of values. Besides we also extracted group “Delayed” (D), those being trees exhibiting mixed value from both groups. Scores of “delayed” clustered above those of the most “Retarded” scores and some measures were distributed across higher ranges, nevertheless their overall performance remained visibly lower than that of the “Normals”.

It is of importance to stress that even though we did call the best plant conditions as “Normal”, we do not definite it is the perfect plant condition. Tree condition, due to perennial phenology, is the result of a series of events that have occurred over the course of numerous growing seasons. That makes morphometry, which is focused on aggregate effects, a good measure to represent the history of the holobiont (Noshita et al., 2022; Hartmann et al., 2021).

The selected morphometric parameters that shown highest potential in assessing growth delays were: AREA_PX, COMPACTNESS, HEIGHT_PX, PERIMETER_PX, WIDTH_PX, ROUNDNESS (Fig. 10). The Pots site shows greater overlap among groups while the Forest site exhibits clearer separation particularly in the side view. Traditional metrics (e.g., branch length, leaf count) followed similar trend, showing significant numerical disparities with three distinct grouping of values (Fig. 10 and Supplementary_3_morphometry_data file). The parameters such as AREA_PX, PERIMETER_PX, quantify the plant’s size and edge complexity and provide valuable insights into plant morphology and growth patterns (Sadeghi-Tehran et al., 2017; Li et al., 2021). This can indicate growth vigour, stress responses, or resource allocation strategies, reflecting underlying biological processes and environmental adaptations (Noshita et al., 2022; Humplík et al., 2015). In our acquired data, differences in those parameters between group “Normal” and “Retarded” which grew in the same site can vary wildly. Depending on the analysed subject differences can reach up to 27 times (AREA_PX) clearly suggesting major influence of external factor. Together, these parameters share similar parallel trend to differences in fungal community structure.

Colour segmentation further validated these groups (Fig. 12). The Normal group exhibited denser, greener leaf mass; the Delayed group had reduced coverage; and the Retarded group showed minimal leaf area and markedly paler crown pigmentation due to higher proportion of branches colours. For Normal group, both in Pots and Forest site, green “class” colours show dominance in percentage compositions indicating higher green-leaf coverage. Paler/tan colours dominance occurs in Retarded and some Delayed seedlings that reflect increased branch exposure consistent with reduced canopy cover.

#### 3.3.3 Test hypothesis validation

##### H1: Different plant conditions are accompanied by different mycobiome structure

Our method was capable of seeing a difference both in plant conditions and accompanying them fungal rhizobiome as described in section 3.3. The difference in mycobiome structure can be summarised as increase of chaos with parallel worsening of plant vitality. It means that seedlings condition show dropping trend parallel to change in fungal community structure. Our method also showed that mycobiome compositions are much more dependent on site than on plant condition.

##### H2: Pots and Forest plants of corresponding condition is accompanied by a similar mycobiome structure

The biggest similarity can be seen in values for species numbers, where the conditions from two sites show similar values with Pots dataset having always a few more species. In terms of guild ratio abundance values, Normal and Delayed conditions do not exhibit similarity with Retarded condition being vaguely similar. Network parameters show biggest similarity for Normal conditions in both sites with less consistent ones for Retarded. Delayed condition was most different between sites.

##### H3: The structure of the mycobiome accompanying the intermediate plant condition has characteristics halfway between those of the best and worst plant condition

Only major similarity between Delayed conditions from two sites is number of species in sample set, but as was said before – two Delayed conditions do not share a majority of species. In terms of trophic composition, the Delayed group is the reason for non-linear nature of the Normal-Delayed-Retarded trend, skewing it. Network parameters show that while they are always an intermediate condition, Delayed in Pots seem to be akin to Retarded and in Forest more akin to Normal condition.

## 4. Conclusions

The proposed GiaC-phenomic methodology was capable of verifying research hypotheses concerning the assessment of the health status of the plant-microbiome system. We proved that GiaC is suitable for the analysis of microbiome. It integrates multiple complementary methods that in tandem provides a more comprehensive characteristic of its state. Interestingly, species taxonomic and trophic composition on its own did not show a trend reflected in plant condition but were best for site difference. Graph based methods were able to show fungal rhizobiome structure similarity between plant conditions from our studied sites but produced specific artifacts if sample sets were mixed. That feature of GiaC’s network analysis allows it to be an implicit sample set homogeneity test.

Unlike conventional phytopathological studies, which predominantly focus on direct fungus-plant interactions (e.g., pathogenic infections or mycorrhizal symbiosis), our approach emphasized on fungus-fungus dynamics within the mycobiome. By integrating low-cost remote sensing morphometric classification as a complementary module to mycobiome profiling, we provide an alternative framework for diagnosing seedlings condition.

This perspective allows us for a more comprehensive understanding of how soil microbial interactions influence seedling health and considers the broader ecological context of microbial communities. Combining GiaC and plant morphometry we were able to observe that more stable fungal co-occurrence patterns should be attributed to the plant set of the best growth. In contrast, for plants with the worse viability, more chaotic patterns can be considered non-optimal for plant-mycobiome cooperation. GiaC was also capable of showing that holobiont of plant in suboptimal, but not critical condition, is not simply a halfway point between optimal and chaotic states, but something entirely separate.

Coincidence of lower plant scores with specific mycobiome structure can be used as early detector or selection tool meant to spot and select disrupted plant-fungal coexistence. By doing so presented method holds promise for applications in widely understood forest restoration, nursery management, and ecological research, particularly in settings where advanced molecular or computational resources are unavailable.

## Supporting information

Morphometry dara

Network visualisations

Sankey flows

## Data availability

Raw data from our sequencing can be found in NCBI SRA database under BioProject PRJNA1261255. Created networks, raw morphometric values, details on guild categorisation of FungalTraits terms and F99 source code can be found in our GitHub repository https://github.com/PunkOkami/GiaC-phenomic_supps

## Acknowledgements

We would like to express our gratitude Prof. Mirosława Dabert, the chief of Molecular Biology Techniques Laboratory at Adam Mickiewicz University, and her team for their expert assistance during the NGS sequencing stage.

We acknowledge Mikołaj Charchuta’s involvement in funding acquisition and DNA isolation and sequencing of soil samples.

We would also like to thank Parv Malik for designing graphical abstract - both the initial draft and well as the final version.

## Funding

The publication was supported by the Polish Ministry of Science and Higher Education of Poland through statutory funds of Adam Mickiewicz University: Department of Plant Physiology (506000/604/4102020105), Department of Environmental Remote Sensing and Soil Science (506000/607/4103010000), AMU students IDUB grant UAM 075/39/ID-UB/0015 for M.Ch., and funds of The National Centre for Research and Development - student internship for M.Ch. under project POWR.03.05.00-00-Z303/17 cofounded by European Social Fund.

## CRediT authorship contribution statement

**Maksymilian Chmielewski**: Methodology, Software, Formal analysis, Investigation, Data Curation, Writing - Original Draft, Writing - Review & Editing, Visualization, Funding acquisition, Validation, Resources; **Jakub Kuncewicz:** Methodology, Formal analysis, Investigation, Writing - Original Draft, Writing - Review & Editing, Visualization Validation, Resources; **Władysław Polcyn**: Conceptualization, Methodology, Investigation, Writing - Original Draft, Writing - Review & Editing, Supervision, Project administration, Funding acquisition, Resources.

## Notes

https://github.com/PunkOkami/GiaC-phenomic_supps

https://www.ncbi.nlm.nih.gov/bioproject/PRJNA1261255

### Competing Interest Statement

The authors have declared no competing interest.

