## Supplementary material for "Multimodal GiaC-phenomic approach for microbiome-tree system profiling": Morphometry dara

### Supplementary image analysis and morphometry data

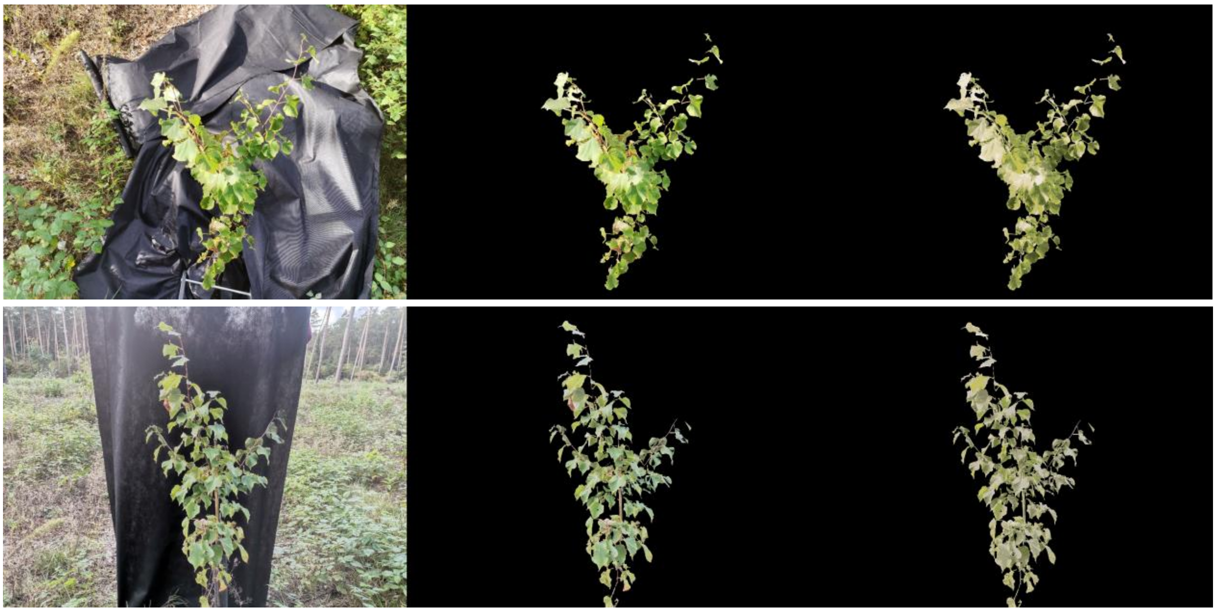

Example of two raw images (left column), their cut out after mask construction (center column) and their colour segmentation (right column). Top panel shows top view and bottom panel shows side views of the crown of tree seedling.

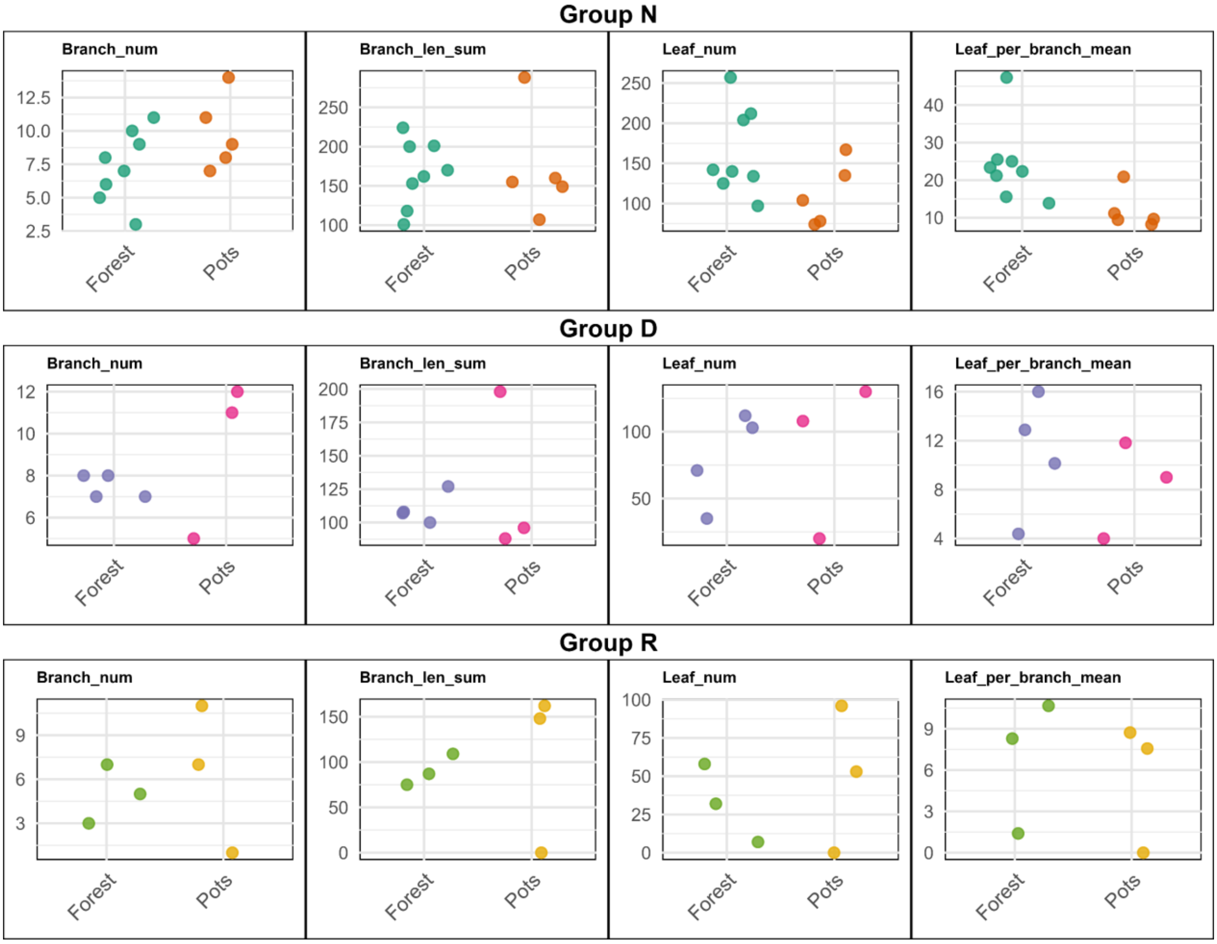

Summary of the variation in manual measurements of branch number (Branch_num) total branch length (Branch_len_sum), leaf count (Leaf_num) and leaf density (Leaf_per_branch_mean) for seedlings grouped by vitality (N = Normal, D = Delayed, R = Retarded) and site (Pots and Forest). Values are plotted on the raw measurement scales (branch length in centimeters, branch number and leaf count as absolute counts, leaf density as the per-plant density metric), with each point representing one seedling observation.

For raw values see our GitHub repository: https://github.com/PunkOkami/GiaC-phenomic_supps
