## Supplementary material for "Multimodal GiaC-phenomic approach for microbiome-tree system profiling": Network visualisations

### Network parameters and ClusterCollapse visualisations

This Supplementary presents ClusterCollapse and Make-up Tables for each of 6 plant conditions except Normal Forest, as that is presented in body text on Figure XYZ. Document also included raw values for all network parameters for all original networks (before ClusterCollapse).

| Parameter | Delayed_forest | Delayed_pots | Normal_forest | Normal_pots | Retarded_forest | Retarded_pots |
|---|---|---|---|---|---|---|
| Edge density | 0.0224 | 0.022 | 0.0221 | 0.0224 | 0.0221 | 0.0232 |
| Species number | 198 | 211 | 175 | 183 | 248 | 251 |
| Cluster number | 13 | 14 | 12 | 10 | 13 | 12 |
| Clustering coeff | 0.0975 | 0.0935 | 0.0888 | 0.0728 | 0.089 | 0.0962 |
| Modularity | 0.5348 | 0.5122 | 0.5593 | 0.5331 | 0.4586 | 0.4436 |
| Natural connectivity | 0.0065 | 0.0062 | 0.0072 | 0.0068 | 0.0054 | 0.0053 |
| Radius | 6 | 5 | 6 | 6 | 5 | 5 |
| Diameter | 10 | 9 | 10 | 12 | 9 | 9 |
| Bridge number | 1 | 3 | 7 | 10 | 0 | 1 |
| Bridge node number | 1 | 3 | 8 | 9 | 0 | 1 |
| Component number | 1 | 1 | 1 | 1 | 1 | 1 |
| Vertex connectivity | 1 | 1 | 1 | 1 | 2 | 1 |
| Edge connectivity | 1 | 1 | 1 | 1 | 2 | 1 |
| Positive edge percentage | 62.844 | 68.2377 | 61.1276 | 59.2493 | 63.6095 | 60.5227 |
| Number of isolated nodes | 0 | 0 | 0 | 0 | 0 | 0 |

Table 1: Raw values of network parameters for all sample sets

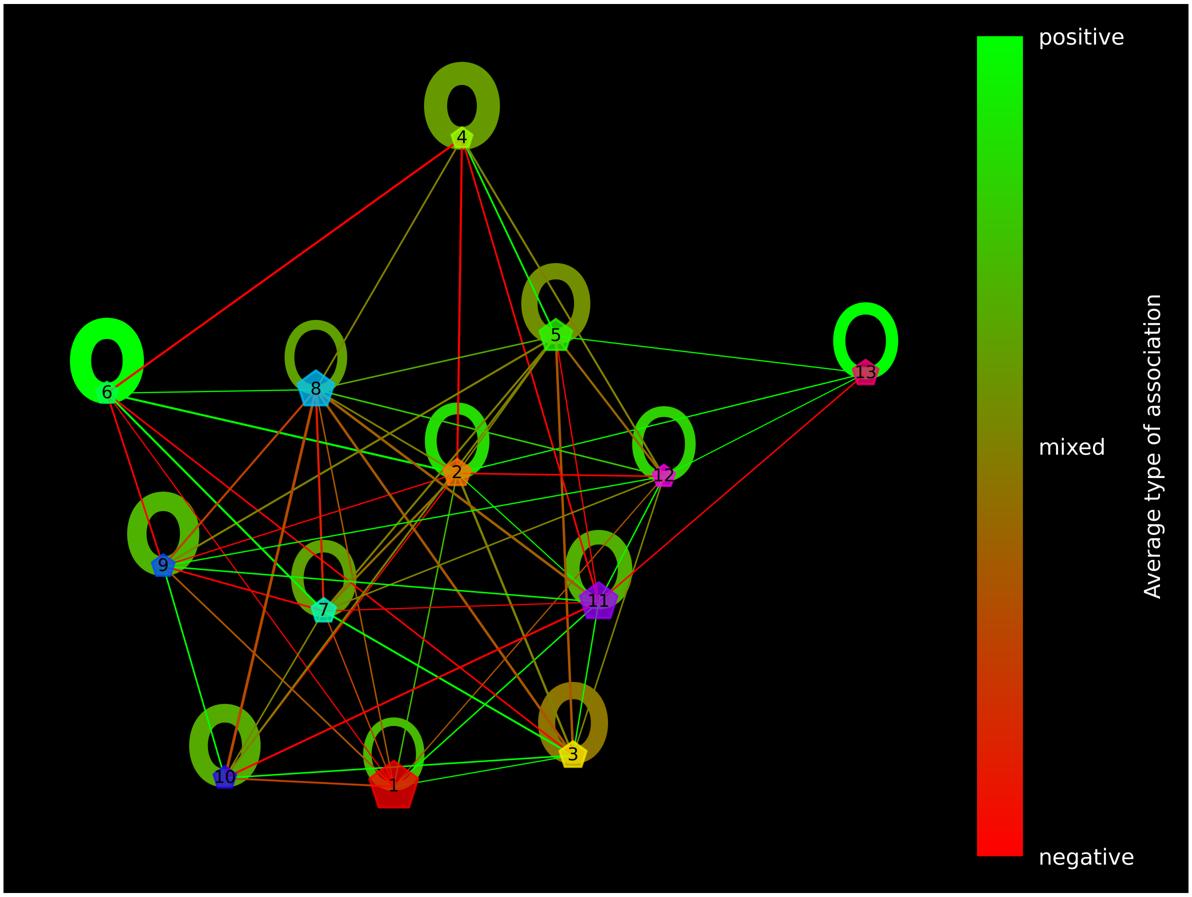


| | ENA | End-Sap | Myc | Npa | Npa-End | Npa-Sap | Pat | Pat-End | Pat-Sap | Sap | Sap-End | Sap-Npa | Sap-Pat |
|---|---|---|---|---|---|---|---|---|---|---|---|---|---|
| Cluster 1 (F: 30.08%, S: 27) | 0.22 | 0.01 | 0.0 | 0.0 | 0.0 | 0.0 | 0.0 | 0.02 | 0.1 | 0.68 | 0.02 | 0.01 | 0.0 |
| Cluster 2 (F: 5.42%, S: 19) | 0.28 | 0.0 | 0.04 | 0.01 | 0.0 | 0.0 | 0.02 | 0.0 | 0.0 | 0.68 | 0.0 | 0.0 | 0.0 |
| Cluster 3 (F: 4.74%, S: 10) | 0.35 | 0.01 | 0.0 | 0.0 | 0.0 | 0.02 | 0.07 | 0.02 | 0.0 | 0.57 | 0.0 | 0.0 | 0.0 |
| Cluster 4 (F: 0.43%, S: 6) | 0.24 | 0.0 | 0.0 | 0.0 | 0.0 | 0.0 | 0.08 | 0.0 | 0.0 | 0.7 | 0.0 | 0.0 | 0.0 |
| Cluster 5 (F: 9.82%, S: 13) | 0.12 | 0.0 | 0.0 | 0.0 | 0.0 | 0.0 | 0.06 | 0.0 | 0.02 | 0.82 | 0.0 | 0.0 | 0.02 |
| Cluster 6 (F: 1.14%, S: 9) | 0.13 | 0.0 | 0.31 | 0.0 | 0.0 | 0.0 | 0.0 | 0.15 | 0.0 | 0.44 | 0.0 | 0.0 | 0.0 |
| Cluster 7 (F: 2.92%, S: 19) | 0.37 | 0.0 | 0.0 | 0.0 | 0.04 | 0.0 | 0.0 | 0.04 | 0.06 | 0.51 | 0.03 | 0.0 | 0.0 |
| Cluster 8 (F: 14.18%, S: 25) | 0.24 | 0.0 | 0.01 | 0.01 | 0.0 | 0.01 | 0.0 | 0.0 | 0.01 | 0.35 | 0.12 | 0.0 | 0.3 |
| Cluster 9 (F: 1.68%, S: 9) | 0.08 | 0.0 | 0.34 | 0.0 | 0.03 | 0.0 | 0.0 | 0.0 | 0.51 | 0.0 | 0.07 | 0.0 | 0.0 |
| Cluster 10 (F: 1.57%, S: 10) | 0.57 | 0.0 | 0.0 | 0.0 | 0.0 | 0.0 | 0.2 | 0.0 | 0.0 | 0.24 | 0.02 | 0.0 | 0.0 |
| Cluster 11 (F: 15.18%, S: 13) | 0.21 | 0.0 | 0.0 | 0.0 | 0.0 | 0.0 | 0.03 | 0.0 | 0.0 | 0.77 | 0.0 | 0.0 | 0.02 |
| Cluster 12 (F: 1.26%, S: 20) | 0.38 | 0.0 | 0.04 | 0.0 | 0.0 | 0.0 | 0.06 | 0.0 | 0.08 | 0.42 | 0.07 | 0.0 | 0.0 |
| Cluster 13 (F: 3.82%, S: 18) | 0.31 | 0.23 | 0.12 | 0.0 | 0.0 | 0.03 | 0.0 | 0.04 | 0.01 | 0.2 | 0.11 | 0.0 | 0.0 |

Figure 1: ClusterCollapse visualisation and Make-up Table for Delayed Forest

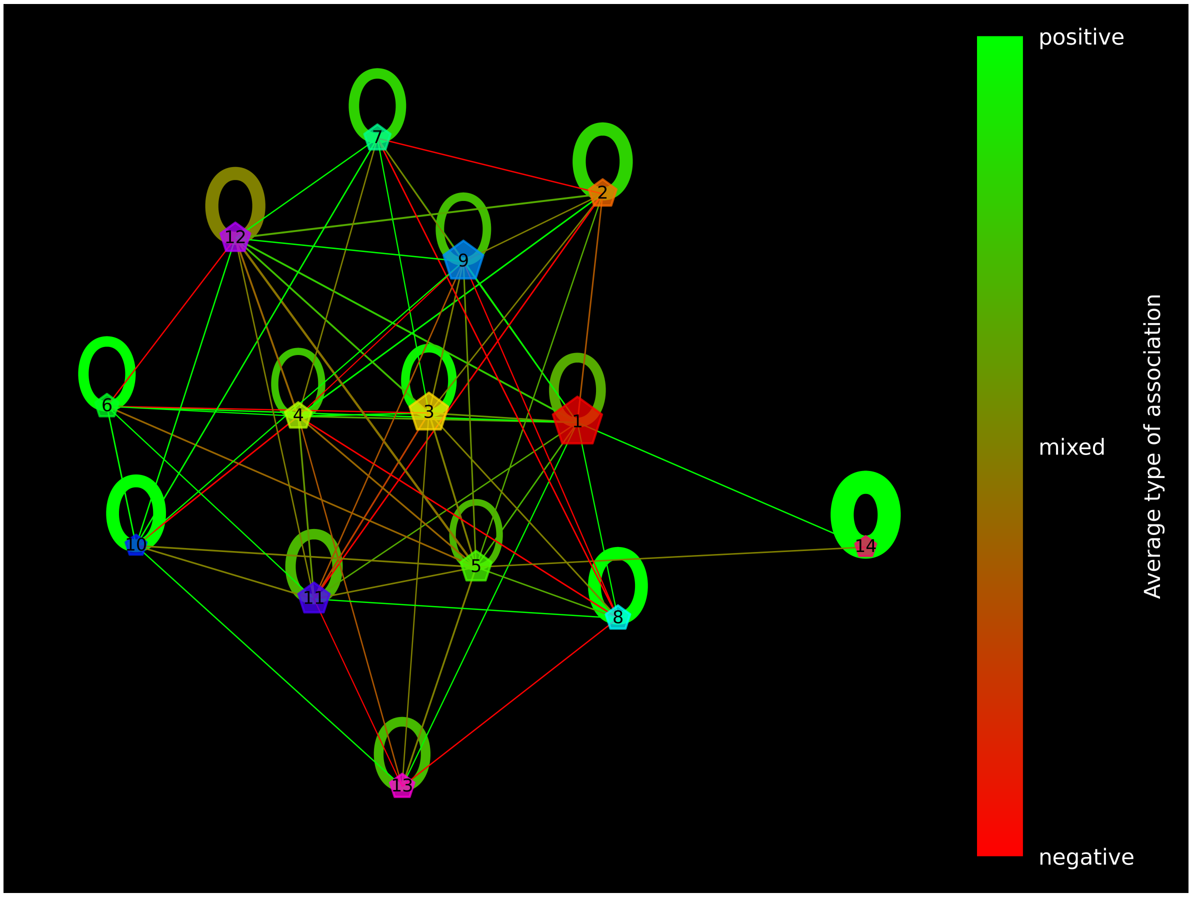


| | ENA | End-Sap | Myc | Npa | Npa-Sap | Pat | Pat-End | Pat-Sap | Sap | Sap-End | Sap-Pat |
|---|---|---|---|---|---|---|---|---|---|---|---|
| Cluster 1 (F: 23.63%, S: 21) | 0.3 | 0.03 | 0.0 | 0.0 | 0.0 | 0.0 | 0.0 | 0.1 | 0.59 | 0.01 | 0.0 |
| Cluster 2 (F: 4.32%, S: 12) | 0.38 | 0.0 | 0.0 | 0.0 | 0.0 | 0.0 | 0.0 | 0.08 | 0.57 | 0.0 | 0.0 |
| Cluster 3 (F: 12.83%, S: 17) | 0.91 | 0.0 | 0.02 | 0.01 | 0.02 | 0.03 | 0.0 | 0.01 | 0.01 | 0.0 | 0.04 |
| Cluster 4 (F: 3.63%, S: 20) | 0.31 | 0.0 | 0.0 | 0.0 | 0.02 | 0.0 | 0.03 | 0.13 | 0.55 | 0.0 | 0.0 |
| Cluster 5 (F: 6.63%, S: 31) | 0.5 | 0.11 | 0.02 | 0.03 | 0.0 | 0.0 | 0.03 | 0.02 | 0.28 | 0.0 | 0.08 |
| Cluster 6 (F: 1.81%, S: 11) | 0.22 | 0.0 | 0.0 | 0.0 | 0.0 | 0.22 | 0.0 | 0.04 | 0.55 | 0.0 | 0.0 |
| Cluster 7 (F: 3.39%, S: 11) | 0.92 | 0.0 | 0.0 | 0.0 | 0.0 | 0.0 | 0.0 | 0.0 | 0.09 | 0.0 | 0.0 |
| Cluster 8 (F: 2.34%, S: 9) | 0.37 | 0.0 | 0.0 | 0.0 | 0.0 | 0.02 | 0.0 | 0.0 | 0.0 | 0.63 | 0.0 |
| Cluster 9 (F: 13.83%, S: 19) | 0.78 | 0.0 | 0.02 | 0.0 | 0.0 | 0.11 | 0.07 | 0.05 | 0.01 | 0.01 | 0.0 |
| Cluster 10 (F: 0.5%, S: 9) | 0.22 | 0.0 | 0.3 | 0.0 | 0.0 | 0.0 | 0.0 | 0.04 | 0.46 | 0.0 | 0.0 |
| Cluster 11 (F: 6.87%, S: 16) | 0.19 | 0.0 | 0.21 | 0.0 | 0.0 | 0.02 | 0.0 | 0.56 | 0.06 | 0.0 | 0.0 |
| Cluster 12 (F: 5.76%, S: 13) | 0.12 | 0.0 | 0.01 | 0.0 | 0.0 | 0.02 | 0.0 | 0.09 | 0.79 | 0.0 | 0.0 |
| Cluster 13 (F: 2.28%, S: 16) | 0.3 | 0.0 | 0.0 | 0.0 | 0.0 | 0.47 | 0.0 | 0.18 | 0.07 | 0.0 | 0.02 |
| Cluster 14 (F: 0.49%, S: 6) | 0.59 | 0.0 | 0.0 | 0.0 | 0.24 | 0.19 | 0.0 | 0.0 | 0.0 | 0.0 | 0.0 |

Figure 2: ClusterCollapse visualisation and Make-up Table for Delayed Pots

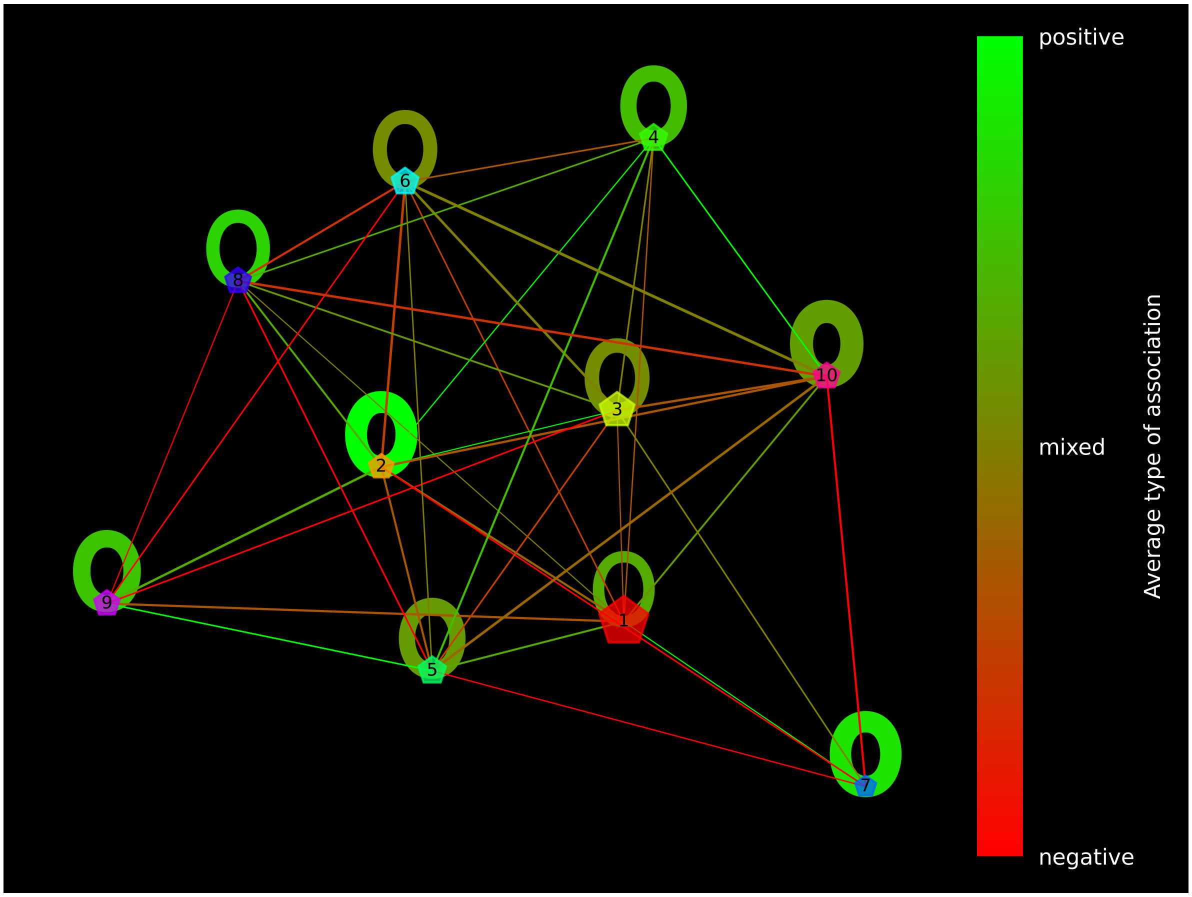


| | ENA | End-Sap | Myc | Npa | Npa-Sap | Pat | Pat-End | Pat-Sap | Sap | Sap-End | Sap-Pat |
|---|---|---|---|---|---|---|---|---|---|---|---|
| Cluster 1 (F: 39.77%, S: 29) | 0.96 | 0.01 | 0.03 | 0.01 | 0.01 | 0.01 | 0.01 | 0.01 | 0.01 | 0.0 | 0.0 |
| Cluster 2 (F: 3.96%, S: 13) | 0.15 | 0.0 | 0.0 | 0.0 | 0.0 | 0.0 | 0.0 | 0.09 | 0.02 | 0.0 | 0.77 |
| Cluster 3 (F: 16.07%, S: 25) | 0.79 | 0.0 | 0.0 | 0.0 | 0.01 | 0.0 | 0.0 | 0.1 | 0.13 | 0.0 | 0.0 |
| Cluster 4 (F: 6.6%, S: 18) | 0.17 | 0.0 | 0.08 | 0.0 | 0.08 | 0.02 | 0.0 | 0.0 | 0.68 | 0.0 | 0.0 |
| Cluster 5 (F: 7.01%, S: 18) | 0.62 | 0.0 | 0.0 | 0.0 | 0.0 | 0.12 | 0.22 | 0.0 | 0.06 | 0.0 | 0.0 |
| Cluster 6 (F: 6.3%, S: 19) | 0.46 | 0.0 | 0.04 | 0.0 | 0.0 | 0.06 | 0.03 | 0.0 | 0.19 | 0.26 | 0.0 |
| Cluster 7 (F: 0.61%, S: 10) | 0.53 | 0.0 | 0.13 | 0.0 | 0.0 | 0.22 | 0.0 | 0.11 | 0.04 | 0.0 | 0.0 |
| Cluster 8 (F: 5.14%, S: 20) | 0.32 | 0.0 | 0.02 | 0.0 | 0.0 | 0.01 | 0.0 | 0.0 | 0.66 | 0.03 | 0.0 |
| Cluster 9 (F: 4.82%, S: 15) | 0.1 | 0.0 | 0.0 | 0.0 | 0.0 | 0.01 | 0.0 | 0.27 | 0.63 | 0.01 | 0.0 |
| Cluster 10 (F: 5.26%, S: 16) | 0.19 | 0.41 | 0.0 | 0.0 | 0.0 | 0.02 | 0.0 | 0.11 | 0.3 | 0.0 | 0.02 |

Figure 3: ClusterCollapse visualisation and Make-up Table for Normal Pots

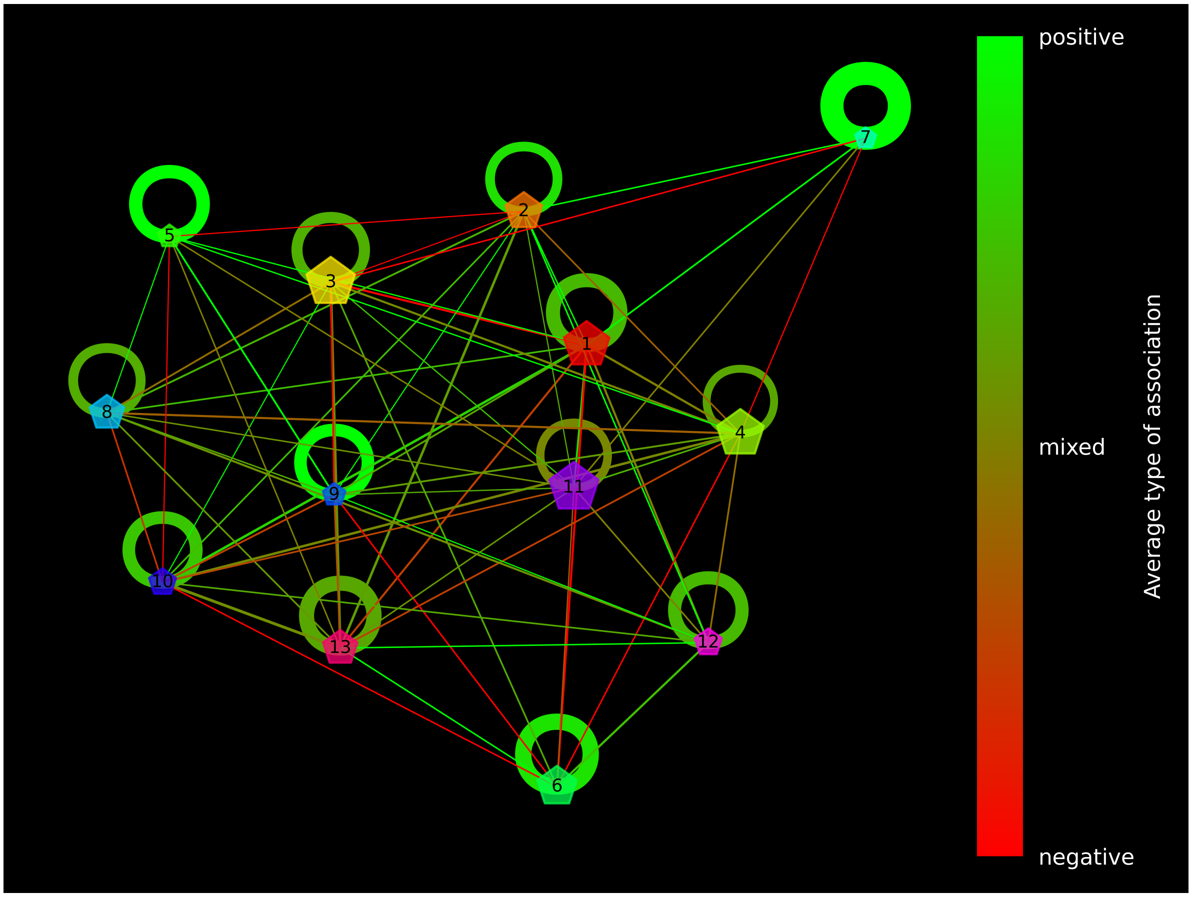


| | ENA | End-Sap | Myc | Npa | Npa-End | Npa-Sap | Pat | Pat-End | Pat-Sap | Sap | Sap-End | Sap-Npa | Sap-Pat |
|---|---|---|---|---|---|---|---|---|---|---|---|---|---|
| Cluster 1 (F: 13.3%, S: 14) | 0.04 | 0.0 | 0.0 | 0.0 | 0.0 | 0.01 | 0.0 | 0.05 | 0.01 | 0.92 | 0.0 | 0.0 | 0.0 |
| Cluster 2 (F: 7.56%, S: 20) | 0.21 | 0.0 | 0.23 | 0.06 | 0.0 | 0.0 | 0.01 | 0.0 | 0.0 | 0.43 | 0.01 | 0.0 | 0.08 |
| Cluster 3 (F: 15.42%, S: 20) | 0.44 | 0.0 | 0.0 | 0.0 | 0.0 | 0.01 | 0.02 | 0.0 | 0.07 | 0.5 | 0.0 | 0.0 | 0.0 |
| Cluster 4 (F: 14.15%, S: 36) | 0.28 | 0.0 | 0.0 | 0.0 | 0.0 | 0.01 | 0.01 | 0.02 | 0.0 | 0.57 | 0.02 | 0.01 | 0.13 |
| Cluster 5 (F: 0.96%, S: 14) | 0.43 | 0.09 | 0.0 | 0.0 | 0.0 | 0.0 | 0.0 | 0.0 | 0.0 | 0.38 | 0.13 | 0.0 | 0.0 |
| Cluster 6 (F: 9.2%, S: 13) | 0.16 | 0.0 | 0.0 | 0.05 | 0.0 | 0.0 | 0.02 | 0.0 | 0.05 | 0.75 | 0.01 | 0.0 | 0.0 |
| Cluster 7 (F: 0.3%, S: 6) | 0.28 | 0.11 | 0.0 | 0.0 | 0.0 | 0.0 | 0.32 | 0.0 | 0.0 | 0.31 | 0.0 | 0.0 | 0.0 |
| Cluster 8 (F: 6.22%, S: 25) | 0.43 | 0.0 | 0.13 | 0.0 | 0.03 | 0.0 | 0.08 | 0.02 | 0.06 | 0.3 | 0.0 | 0.0 | 0.0 |
| Cluster 9 (F: 0.89%, S: 16) | 0.41 | 0.0 | 0.04 | 0.0 | 0.05 | 0.04 | 0.0 | 0.0 | 0.07 | 0.35 | 0.04 | 0.06 | 0.0 |
| Cluster 10 (F: 2.59%, S: 18) | 0.48 | 0.0 | 0.0 | 0.0 | 0.0 | 0.0 | 0.01 | 0.1 | 0.11 | 0.29 | 0.05 | 0.0 | 0.0 |
| Cluster 11 (F: 15.91%, S: 33) | 0.22 | 0.02 | 0.01 | 0.01 | 0.0 | 0.0 | 0.06 | 0.07 | 0.01 | 0.63 | 0.04 | 0.0 | 0.0 |
| Cluster 12 (F: 2.66%, S: 16) | 0.1 | 0.0 | 0.14 | 0.0 | 0.0 | 0.0 | 0.35 | 0.14 | 0.0 | 0.31 | 0.0 | 0.0 | 0.0 |
| Cluster 13 (F: 6.25%, S: 17) | 0.35 | 0.0 | 0.04 | 0.0 | 0.0 | 0.0 | 0.0 | 0.0 | 0.0 | 0.64 | 0.0 | 0.0 | 0.0 |

Figure 4: ClusterCollapse visualisation and Make-up Table for Retarded Forest

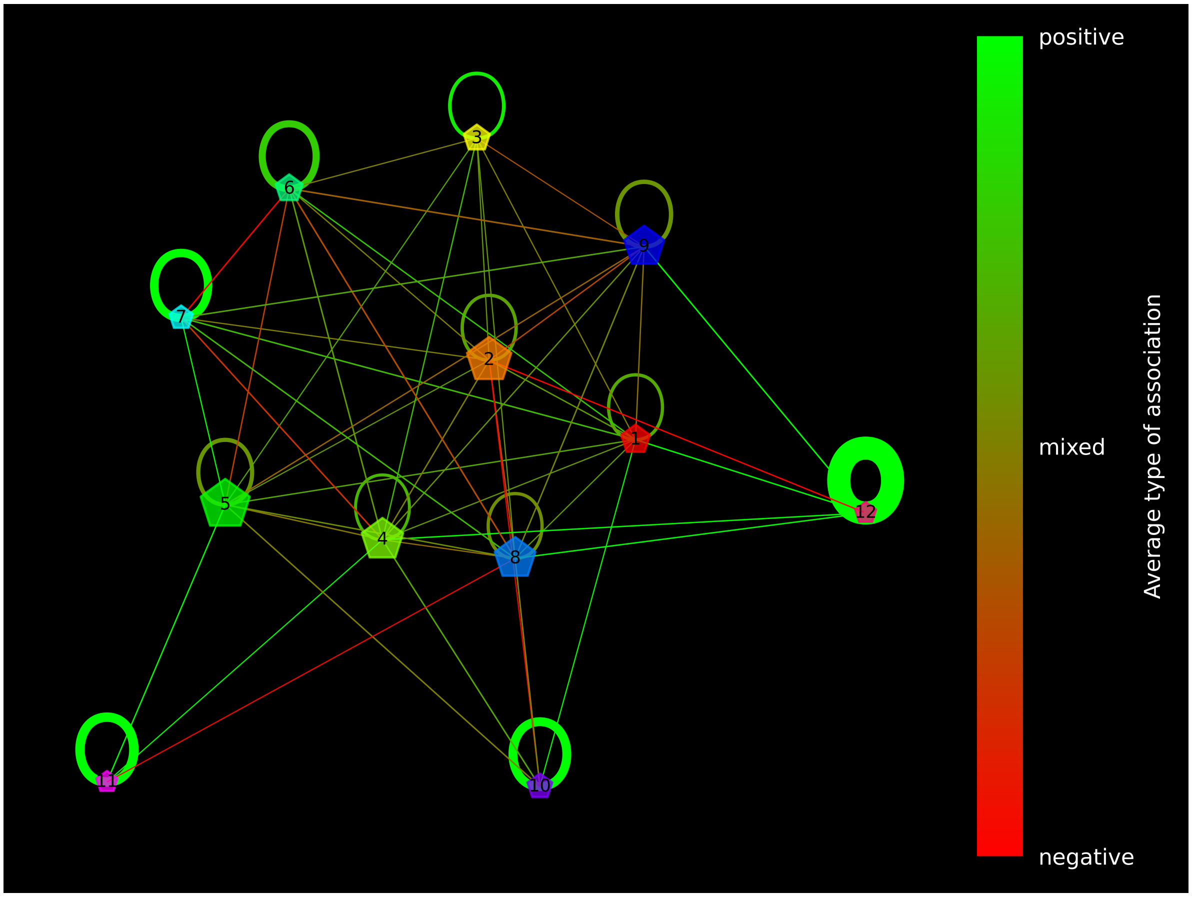


| | ENA | End-Sap | Myc | Npa | Npa-Sap | Pat | Pat-End | Pat-Npa | Pat-Sap | Sap | Sap-End | Sap-Pat |
|---|---|---|---|---|---|---|---|---|---|---|---|---|
| Cluster 1 (F: 4.51%, S: 33) | 0.67 | 0.0 | 0.04 | 0.0 | 0.13 | 0.01 | 0.0 | 0.0 | 0.02 | 0.12 | 0.04 | 0.0 |
| Cluster 2 (F: 17.39%, S: 34) | 0.18 | 0.0 | 0.0 | 0.02 | 0.0 | 0.01 | 0.05 | 0.0 | 0.02 | 0.76 | 0.0 | 0.0 |
| Cluster 3 (F: 2.83%, S: 21) | 0.15 | 0.05 | 0.0 | 0.0 | 0.0 | 0.0 | 0.0 | 0.0 | 0.64 | 0.16 | 0.0 | 0.04 |
| Cluster 4 (F: 14.55%, S: 38) | 0.16 | 0.11 | 0.02 | 0.0 | 0.01 | 0.38 | 0.0 | 0.01 | 0.11 | 0.21 | 0.05 | 0.02 |
| Cluster 5 (F: 22.34%, S: 26) | 0.35 | 0.0 | 0.06 | 0.0 | 0.0 | 0.01 | 0.0 | 0.0 | 0.01 | 0.49 | 0.12 | 0.02 |
| Cluster 6 (F: 3.44%, S: 11) | 0.18 | 0.0 | 0.0 | 0.0 | 0.1 | 0.02 | 0.0 | 0.0 | 0.0 | 0.72 | 0.0 | 0.0 |
| Cluster 7 (F: 1.4%, S: 6) | 0.1 | 0.0 | 0.0 | 0.03 | 0.0 | 0.0 | 0.0 | 0.0 | 0.0 | 0.9 | 0.0 | 0.0 |
| Cluster 8 (F: 13.88%, S: 37) | 0.33 | 0.0 | 0.0 | 0.0 | 0.34 | 0.03 | 0.04 | 0.0 | 0.08 | 0.21 | 0.0 | 0.0 |
| Cluster 9 (F: 13.22%, S: 28) | 0.33 | 0.35 | 0.0 | 0.0 | 0.0 | 0.1 | 0.02 | 0.0 | 0.0 | 0.23 | 0.0 | 0.0 |
| Cluster 10 (F: 2.13%, S: 8) | 0.04 | 0.0 | 0.0 | 0.0 | 0.0 | 0.02 | 0.01 | 0.0 | 0.94 | 0.02 | 0.0 | 0.0 |
| Cluster 11 (F: 0.18%, S: 7) | 1.0 | 0.0 | 0.0 | 0.0 | 0.0 | 0.0 | 0.0 | 0.0 | 0.0 | 0.0 | 0.0 | 0.0 |
| Cluster 12 (F: 0.09%, S: 2) | 0.67 | 0.0 | 0.0 | 0.0 | 0.0 | 0.0 | 0.0 | 0.0 | 0.0 | 0.0 | 0.34 | 0.0 |

Figure 5: ClusterCollapse visualisation and Make-up Table for Retarded Pots
