## Supplementary material for "Multimodal GiaC-phenomic approach for microbiome-tree system profiling": Sankey flows

### Sankey diagrams

This Appendix presents values for number of ASVs for Forest and Pots sample sets. Here we also present number of reads on each filtering step for all 6 plant conditions separately

#### 1 ASV Sankey flow

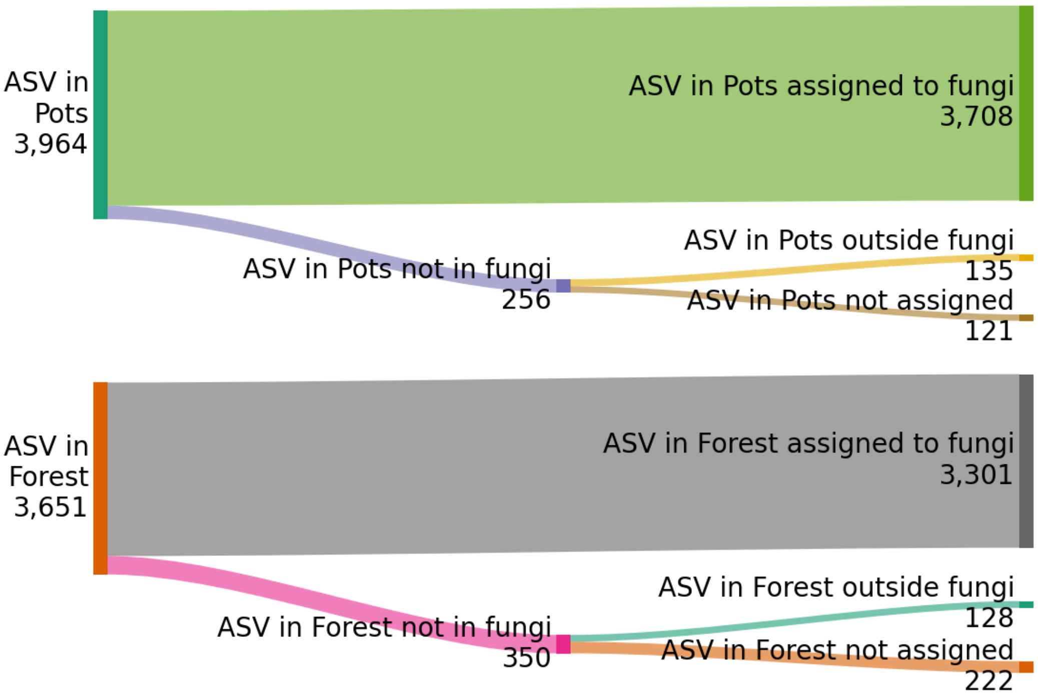


Figure 1: ASV Sankey flow for Forest and Pot sample sets

#### 2 Full read flows for each of 6 plant condition variants

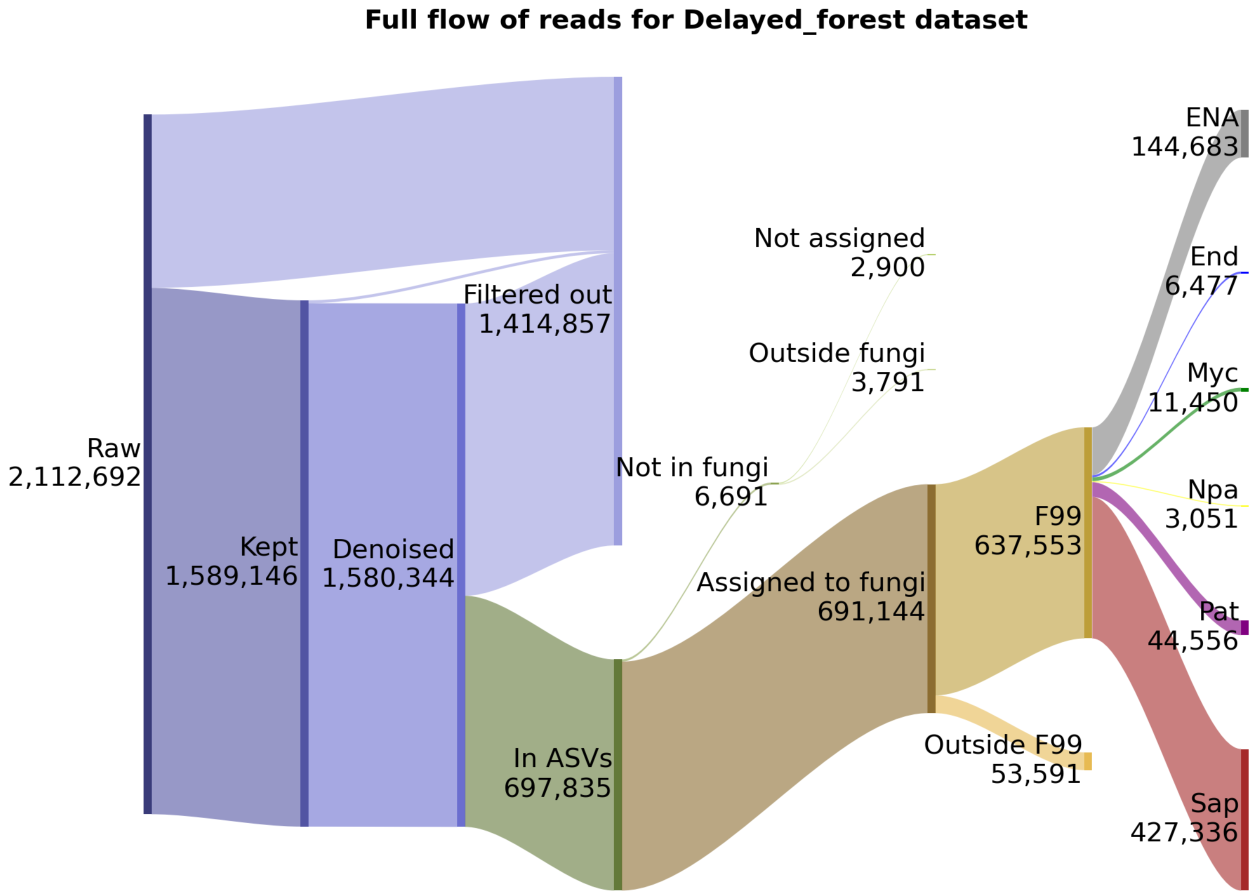


Figure 2: Flow of reads on each filtering step for Delayed Forest

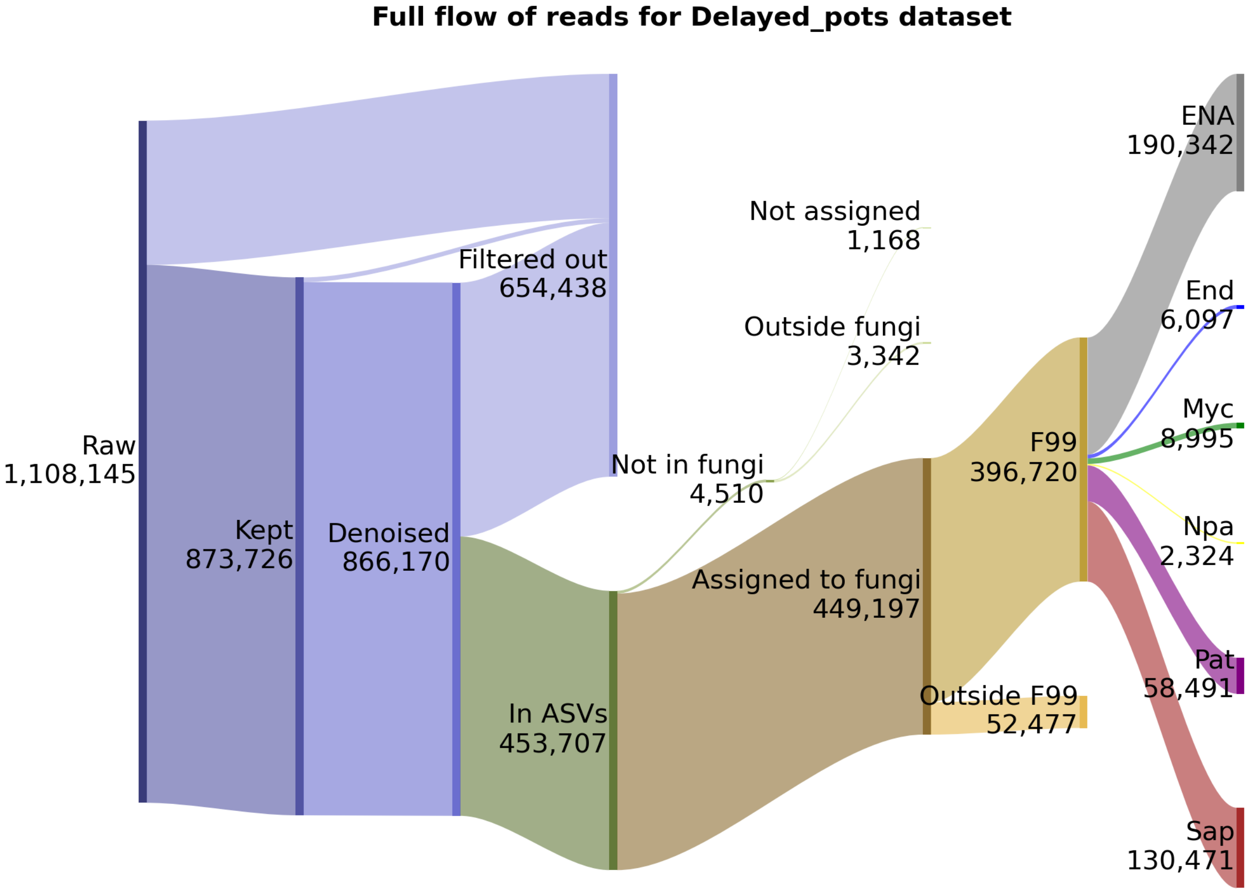


Figure 3: Flow of reads on each filtering step for Delayed Pots

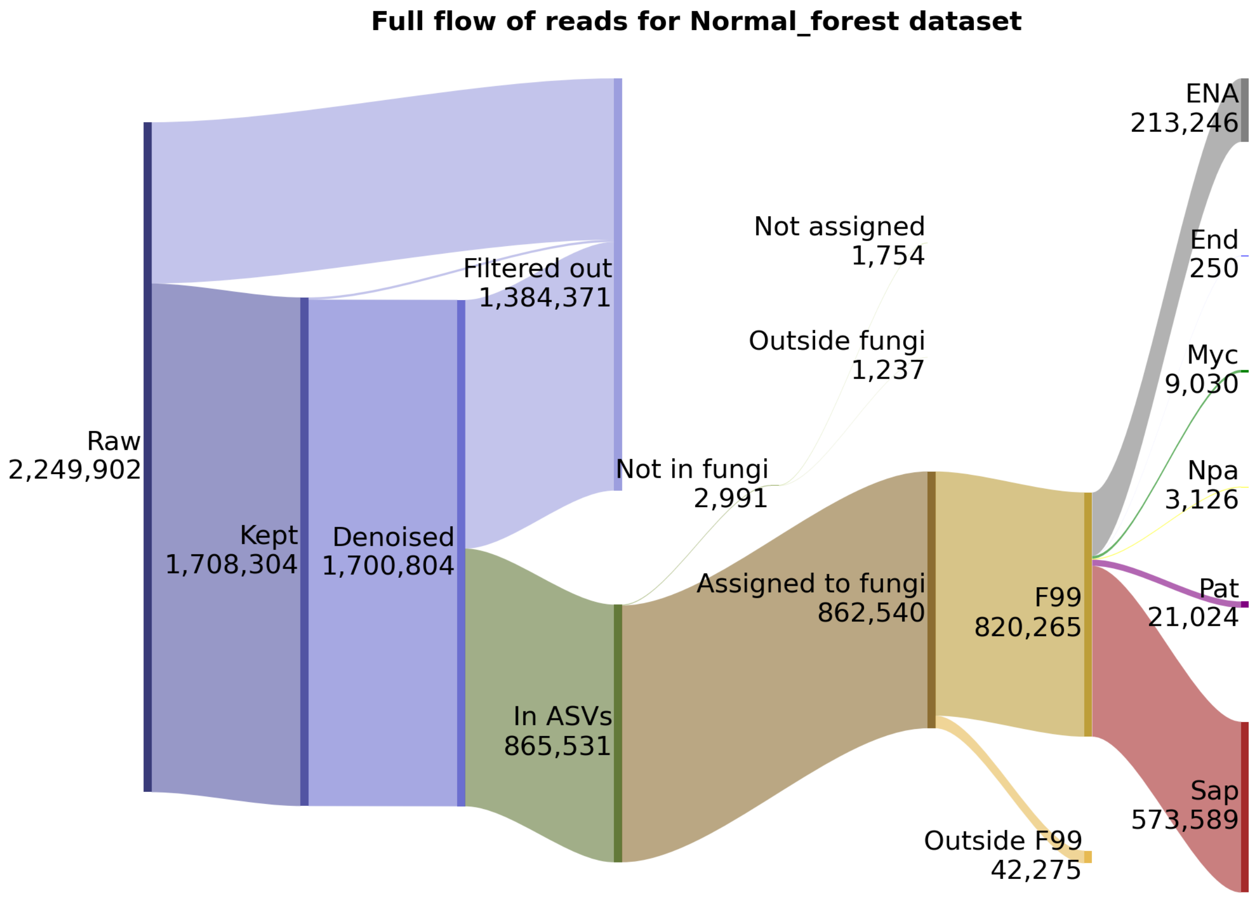


Figure 4: Flow of reads on each filtering step for Normal Forest

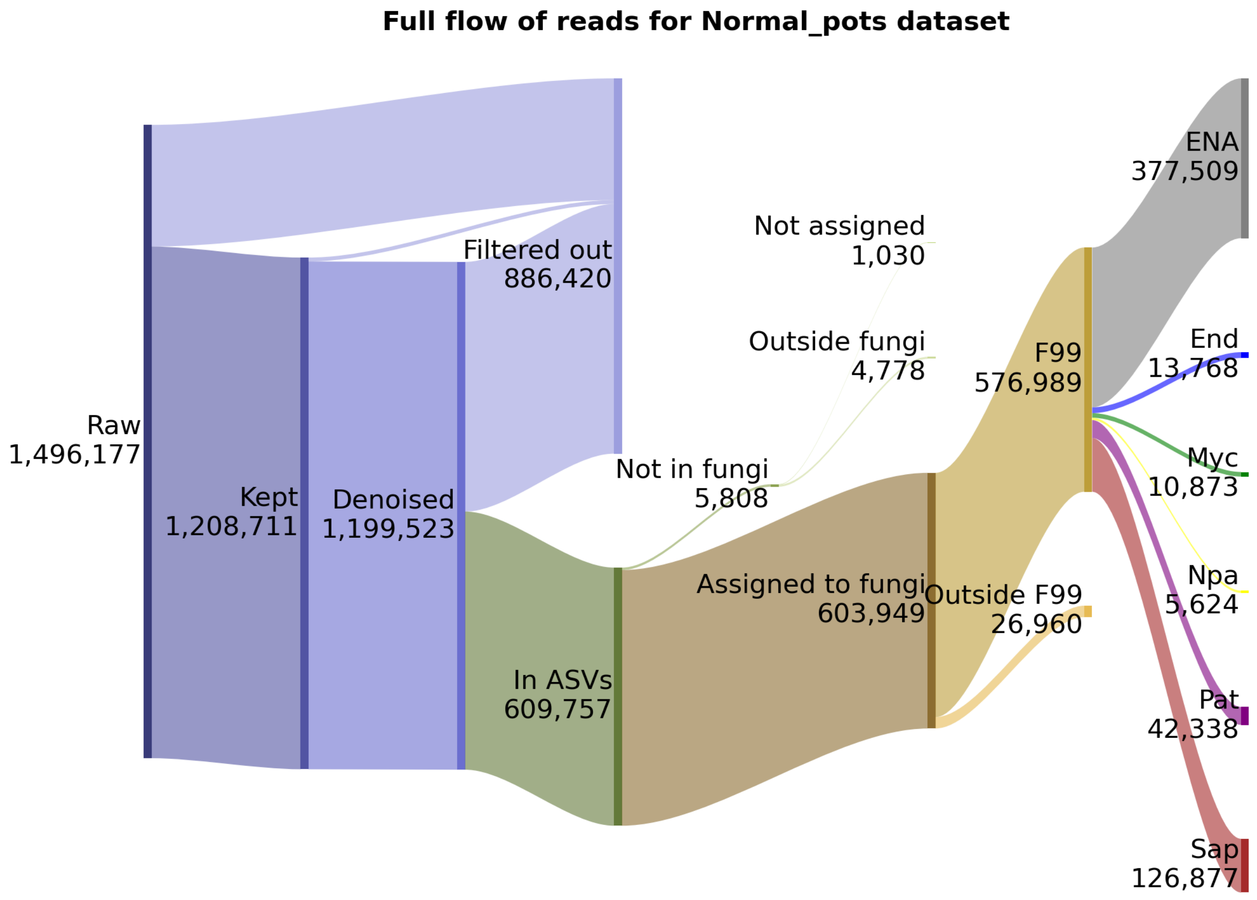


Figure 5: Flow of reads on each filtering step for Normal Pots

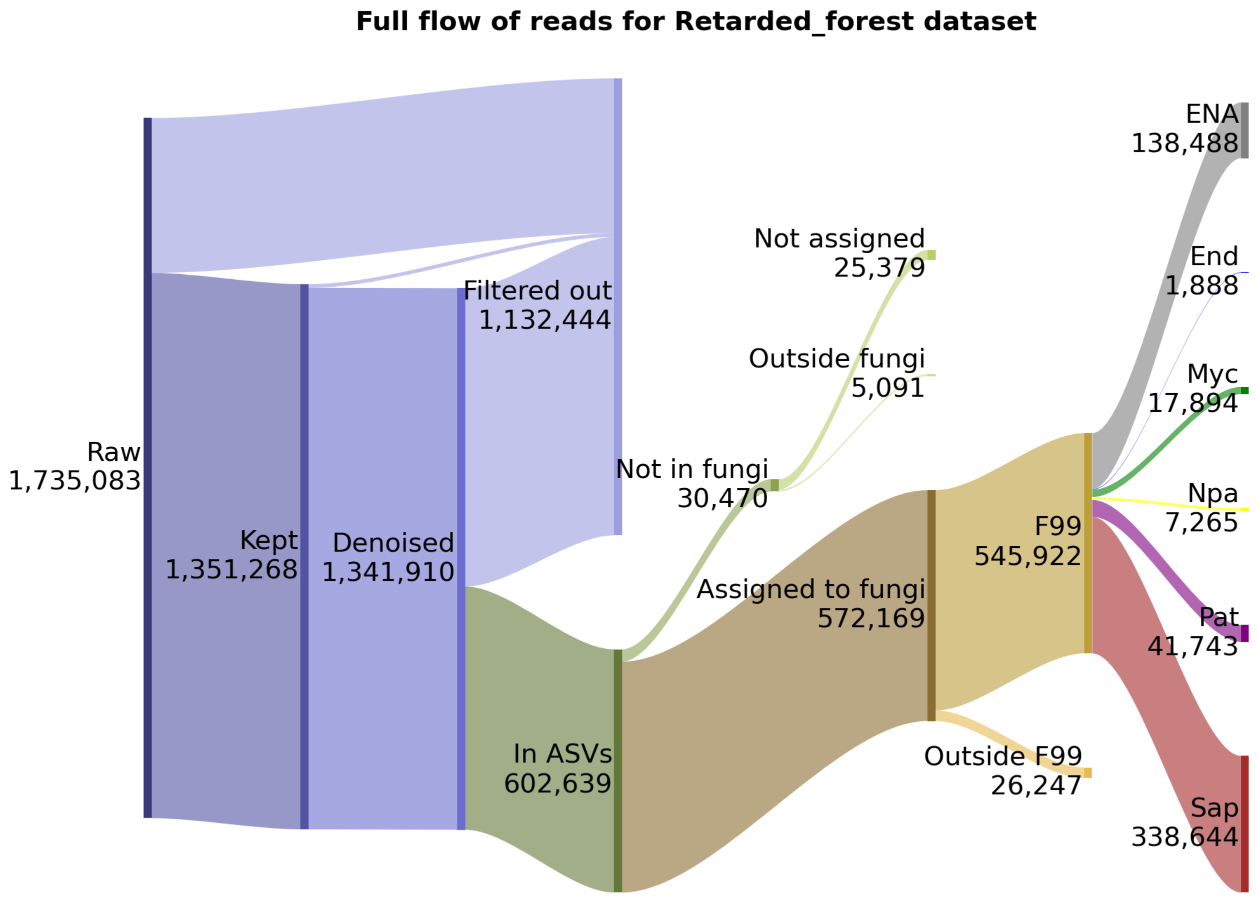


Figure 6: Flow of reads on each filtering step for Retarded Forest

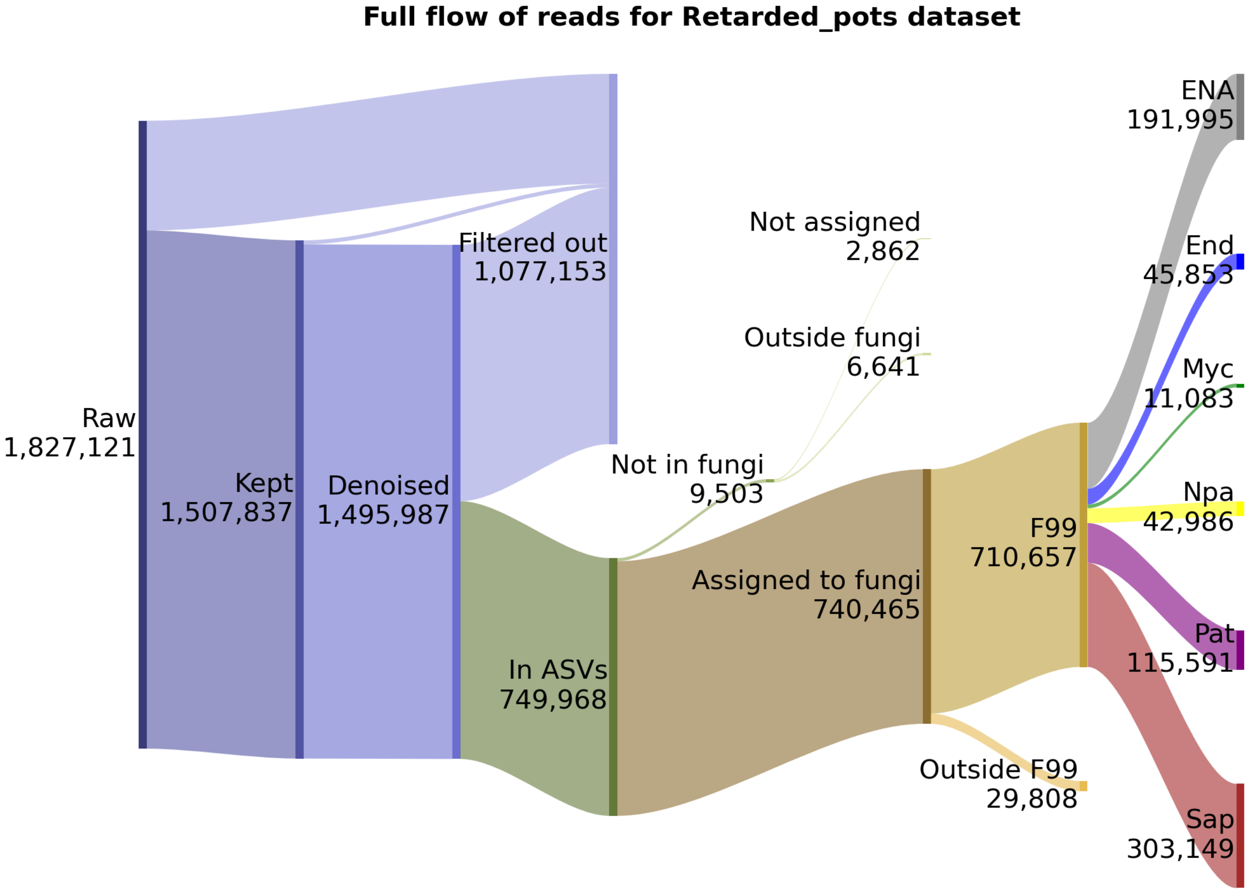


Figure 7: Flow of reads on each filtering step for Retarded Pots
